# Direct androgen receptor regulation of sexually dimorphic gene expression in the mammalian kidney

**DOI:** 10.1101/2023.05.06.539585

**Authors:** Lingyun Xiong, Jing Liu, Seung Yub Han, Kari Koppitch, Jin-Jin Guo, Megan Rommelfanger, Fan Gao, Ingileif B. Hallgrimsdottir, Lior Pachter, Junhyong Kim, Adam L. MacLean, Andrew P. McMahon

## Abstract

Mammalian organs exhibit distinct physiology, disease susceptibility and injury responses between the sexes. In the mouse kidney, sexually dimorphic gene activity maps predominantly to proximal tubule (PT) segments. Bulk RNA-seq data demonstrated sex differences were established from 4 and 8 weeks after birth under gonadal control. Hormone injection studies and genetic removal of androgen and estrogen receptors demonstrated androgen receptor (AR) mediated regulation of gene activity in PT cells as the regulatory mechanism. Interestingly, caloric restriction feminizes the male kidney. Single-nuclear multiomic analysis identified putative cis-regulatory regions and cooperating factors mediating PT responses to AR activity in the mouse kidney. In the human kidney, a limited set of genes showed conserved sex-linked regulation while analysis of the mouse liver underscored organ-specific differences in the regulation of sexually dimorphic gene expression. These findings raise interesting questions on the evolution, physiological significance, and disease and metabolic linkage, of sexually dimorphic gene activity.

## Introduction

Increasing evidence points to differences in epidemiology, pathophysiology, drug responsiveness and disease outcomes between the sexes. For example, human kidney studies indicate age-related decline in renal function is faster in men than in age-matched premenopausal women^1^. Further, chronic disease tends to be more aggressive in men and progresses to end-stage renal disease more rapidly than in women^1, 2^. Men are more susceptible to acute kidney injury, while women are resilient and show improved tolerance to renal ischemia^3–5^. Sex-dependent response to kidney diseases has also been reported in rodents^5–7^. An improved understanding of cellular roles and molecular controls in the male and female kidney will advance our understanding of renal function and renal disease mechanisms between the sexes.

In the mouse, the most widely studied mammalian model, researchers have identified differences in cellular morphology^8–11^, renal physiology^12–14^, and cell type specific gene expression^15–18^ over the last three decades. More recently, a detailed single-cell analysis of gene expression throughout the adult mouse kidney identified 984 genes with sex-biased expression, highlighting proximal tubule (PT) segments as the predominant cellular source of sex-specific variability in gene expression^19^ and microdissection and multiomics approaches identified sex differences in transcription, chromatin accessibility and proteinomics^20^. Proximal tubule cell types play a central role in renal physiology responsible for the primary resorption and recovery of essential molecules from the initial renal filtrate, including glucose, salts and water, and a variety of other important cellular functions such as gluconeogenesis and molecular detoxification^21, 22^.

Sex hormones have long been associated with sex differences in the structure and function of the kidney^23–25^. Androgens enhance salt reabsorption^26^ and water handling^27^ in the PT and stimulate total kidney volume in males^26, 28^. Testosterone has also been shown to modulate urinary calcium clearance^29^, as well as ammonia metabolism and excretion^30, 31^. Gonadal removal and hormone injection studies point to a role for testosterone regulation of sexual dimorphism in both the mouse and rat kidney^23, 30–38^. However, the direct actions of sex hormones and their receptors in regulating sex-biased transcription in the PT has not been studied. Regulatory control of sex-dependent gene expression in the liver has revealed that sex hormones act at the level of the hypothalamic-pituitary axis to control the release of growth hormone (GH), the direct regulator of sexual dimorphic gene expression in hepatocytes^39–44^.

In this study, we investigated the temporal, spatial and genomic regulation of sex hormone action in the mouse kidney. In contrast to the liver, testosterone is the primary and direct driver of sexual dimorphism, acting through Ar receptor regulation of chromatin accessibility in PT cell-types. Complementary genetic studies in the liver revealed a hitherto unrecognized component of direct Ar action on hepatocyte gene expression, with a conserved sex bias in expression of shared genes with the mouse kidney. Comparing gene expression in the mouse and human kidney identified few non-sex chromosome-linked sex-biased genes between the sexes but a conserved sex bias was observed in their expression.

## Results

### Gene-and transcript-level renal sex differences in adult mice

To identify a set of sex-biased genes that are invariant with respect to age, mouse strain and technology assessing gene activity (**Fig. 1A**), we first examined differential gene expression between male and female kidneys in 8-week C57BL/6 mice using whole-kidney bulk RNA-seq (STAR Methods), identifying 1,733 genes with sex-biased expression: 869 expressed at higher levels in the female kidney (female [F]-biased) and 864 expressed at higher levels in the male kidney (male [M]-biased; Table S1.1; *the full set*). We compared this gene set with renal sex differences identified among genetically diverse mice at 6, 12, and 18 months^45^ (GSE121330, referred to as the *JAX* data hereafter) identifying 214 F-biased genes (25%) and 337 M-biased genes (39%) with consistent sex bias across all whole-kidney bulk RNA-seq datasets (551 genes in total; **Fig. 1B**; Table S1.2). This core set of sex-biased genes overlapped significantly with sex-biased gene expression identified in PT segments from single-cell RNA-seq analysis^19^ (PT-sex genes; hypergeometric test, p = 6.0E-27 in female, p = 6.5E-39 in male; **Fig. S1A**). Moreover, the core set includes 78-79% of the PT-sex genes that exhibited persistent sex bias from 6 to 18 months (**Fig. 1B**), including several transcriptional regulators such as *Foxq1* (F-biased) and *Nr1h4* (M-biased).

**Figure 1.**
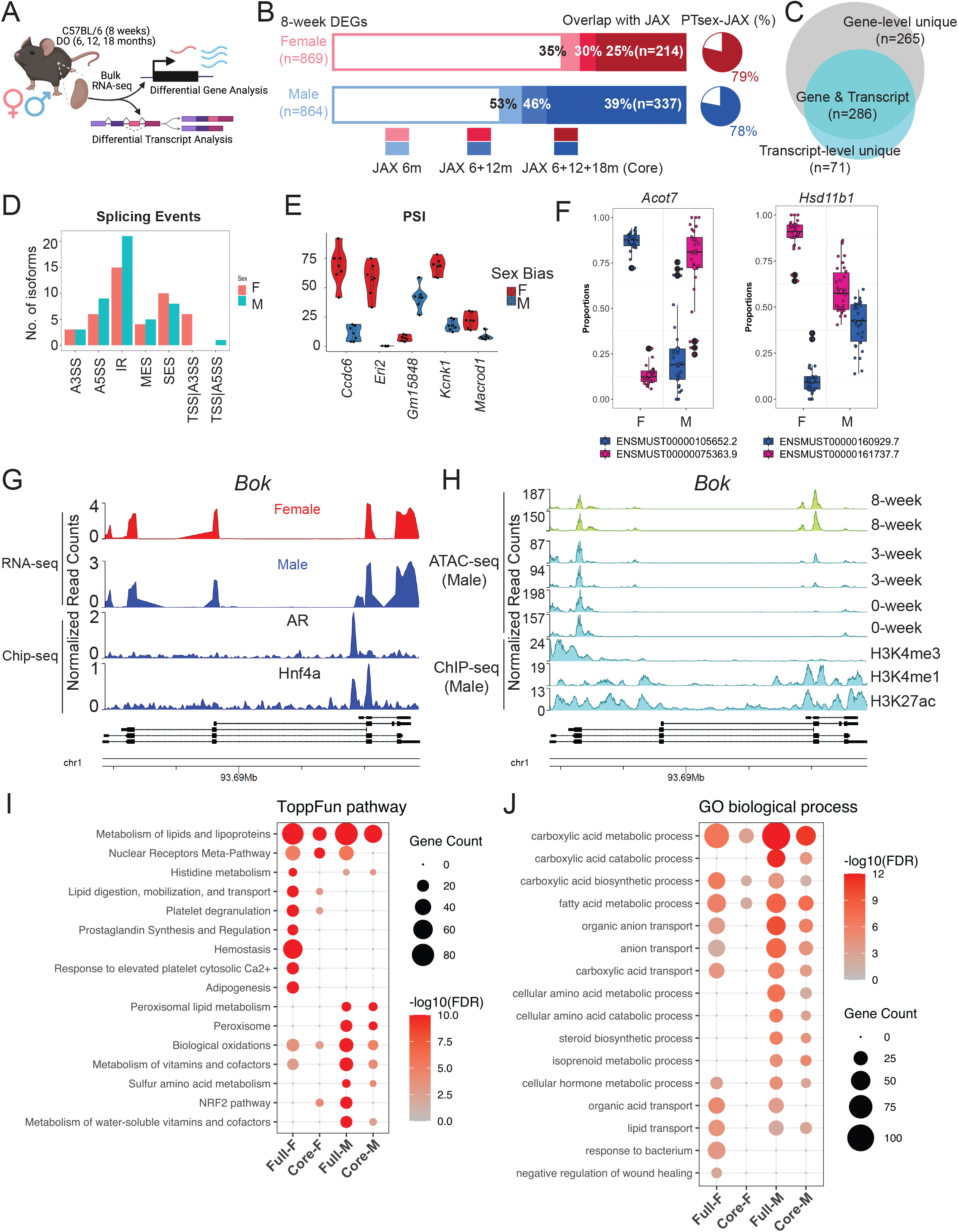
Gene-and transcript-level renal sex differences in adult mice. (A) Schematic summary of the computational analyses of renal transcriptome in adult male and female mice. (B) Stacked bar plot depicts the proportion of sex-biased genes identified in 8-week C57BL/6 mice showing consistent biases in the aging kidney in the diversity outbred (DO) mice (the JAX data^45^). Pie charts represent the percentage of core sex-biased genes that were identified in the PT segments from previous scRNA-seq experiment^19^. (C) Venn diagram compares renal sex differences revealed by gene-and transcript-level analysis. (D) Bar plot shows the distribution of dimorphic isoforms with alternative splicing events in the male and female kidneys. (E) Distribution of Percent Spliced-In values for top genes showing dimorphic splicing events in the male and female kidneys. (F) Dimorphic transcript usage of distinct isoform in *Acot7* (F-biased) and *Hsd11b1* (M-biased) in male and female kidneys. (G-H) Coverage plot over the genomic region of *Bok* by bulk RNA-seq in male and female kidneys, aligned with data from ChIP-seq experiment against AR and Hnf4a in the male kidneys^46^ (G) and by ATAC-seq experiment in the male kidneys^48^, aligned with ENCODE data from ChIP-seq experiment against epigenetic biomarkers^47^ (H). (I-J) Dot plot shows the enrichment of ToppFun pathways (I) and Gene Ontology terms (J) for both the full and core set of sex-biased genes.

At the transcript level, we defined a core set of sex-biased isoforms using the same criteria (**Fig. S1B**). Among the 551 sex-biased genes in the core set, 286 (52%) also showed sex bias in transcript usage (**Fig. 1C**). Interestingly, 71 genes that did not exhibit sex difference at the level of overall gene expression were found to show sex difference at the transcript level (**Fig. 1C**; Table S1.3). Transcript-level sex differences manifest in a variety of alternative splicing events (**Fig. 1D**), including alternative 5’ splice site usage (*Ccdc6*, *Kcnk1* and Macrod1) and intron retention (*Eric* and *Gm15848*) (**Fig. 1E; Fig. S1C**). Among the 286 genes exhibiting sex differences at gene-and transcript-level, *Acot7* (F-biased) and *Hsd11b1* (M-biased) showed the largest disparity in transcript usage (**Fig. 1F; Fig. S1D**). In addition, adult male kidneys express a short isoform of *Bok* (encoding BCL2 Family Apoptosis Regulator) through alternative promoter usage. Comparison of *Bok* transcripts with published studies of Ar chromatin association in the male kidney^46^ and epigenomic histone modifications (H3K4me1 and H3K27ac^47^) in adult male kidneys (**Fig. 1G-H**) indicate the male-specific short isoform of *Bok* has proximal Ar binding associated with chromatin opening in the maturing postnatal kidney^48^ (**Fig. 1H**).

Functional enrichment analysis based on our sex-biased genes and isoforms provided insights into the dimorphic functionalities in the kidney. Using ToppCluster^49^, both sexes showed a significant enrichment in pathways associated with metabolism and lipid lipoproteins, males showed a strong bias in peroxisome lipid metabolism (**Fig. 1I; Fig. S1E**). The male kidney exhibits enhanced expression of the fatty acid translocase Cd36^50^ and acyl-CoA oxidase 1 (encoded by *Acox1*) which catalyzes the first step in peroxisomal fatty acid degradation^51^ while the female kidney shows elevated expression of genes involved in lipid synthesis (*Scd1*)^52^, lipid digestion and mobilization (*Fabp1*)^53^, and the prevention of lipotoxicity (*Acot7*)^54^. Nuclear receptors (NR), which play important roles in maintaining renal function^55^, show differential enrichment in sex-biased gene expression. *Nr1h4*, encoding the farnesoid X nuclear receptor (FXR), is associated with the metabolic shift from synthesis to the oxidation and catabolism of lipids and exhibits a M-bias.

In contrast, expression of several nuclear receptors associated with xenobiotic metabolism showed a F-bias: *Nr1h3*, encoding liver X receptor (LXR); *Nr1i2,* encoding pregnane X receptor (PXR); *Nr1i3*, encoding constitutive androstane receptor (CAR); and *Rxrg, encoding the* retinoid X Receptor gamma sub-unit (RXRγ). In addition, gene ontology (GO) terms for carboxylic acid degradation, amino acid metabolism and steroid biosynthesis were more strongly associated with male kidneys, while female kidneys showed GO term bias associated with bacteria response and negative regulation of wound healing (**Fig. 1J**).

### Male and female renal transcriptomes diverge at puberty between 3-8 weeks

To understand how dimorphic gene expression arises in the mouse kidney^56^, we performed bulk RNA-seq analysis of C57BL/6 male and female kidneys at 0, 2, 4, 8 and 79 weeks post-partum and identified differences in mRNA transcripts between the sexes (**Fig. 2A**). Principle component analysis (PCA) highlighted age (PC1) and sex (PC2) as the leading components of variation in gene expression amongst the kidney samples; sex differences became evident from 4 weeks (**Fig. 2B**). Differential gene expression between male and female kidneys was assessed at each timepoint (**Fig. 2C**; Table S1). In the newborn and 2 week kidney, only sex chromosome encoded genes distinguished the two sexes: 0 and 2-week F-bias in expression of X-linked *Xist* gene and 2 week M-bias in expression of Y-linked genes *Ddx3y, Eif2s3y, Uty*, and *Kdm5d* (Table S1.4-5). In contrast, a pronounced sex bias was observed in a large number of autosomal encoded genes at 4 weeks as mice entered puberty (457 of the 467 genes [97.6%] displaying sex-bias; Table S1.6), which was further enhanced in the kidney at sexual maturity (8 weeks; 1680 of 1733 genes [96.9%] displaying sex bias; Table S1.1) and late life (79 weeks; 1504 of genes [96.7%] displaying sex bias; Table S1.7).

**Figure 2.**
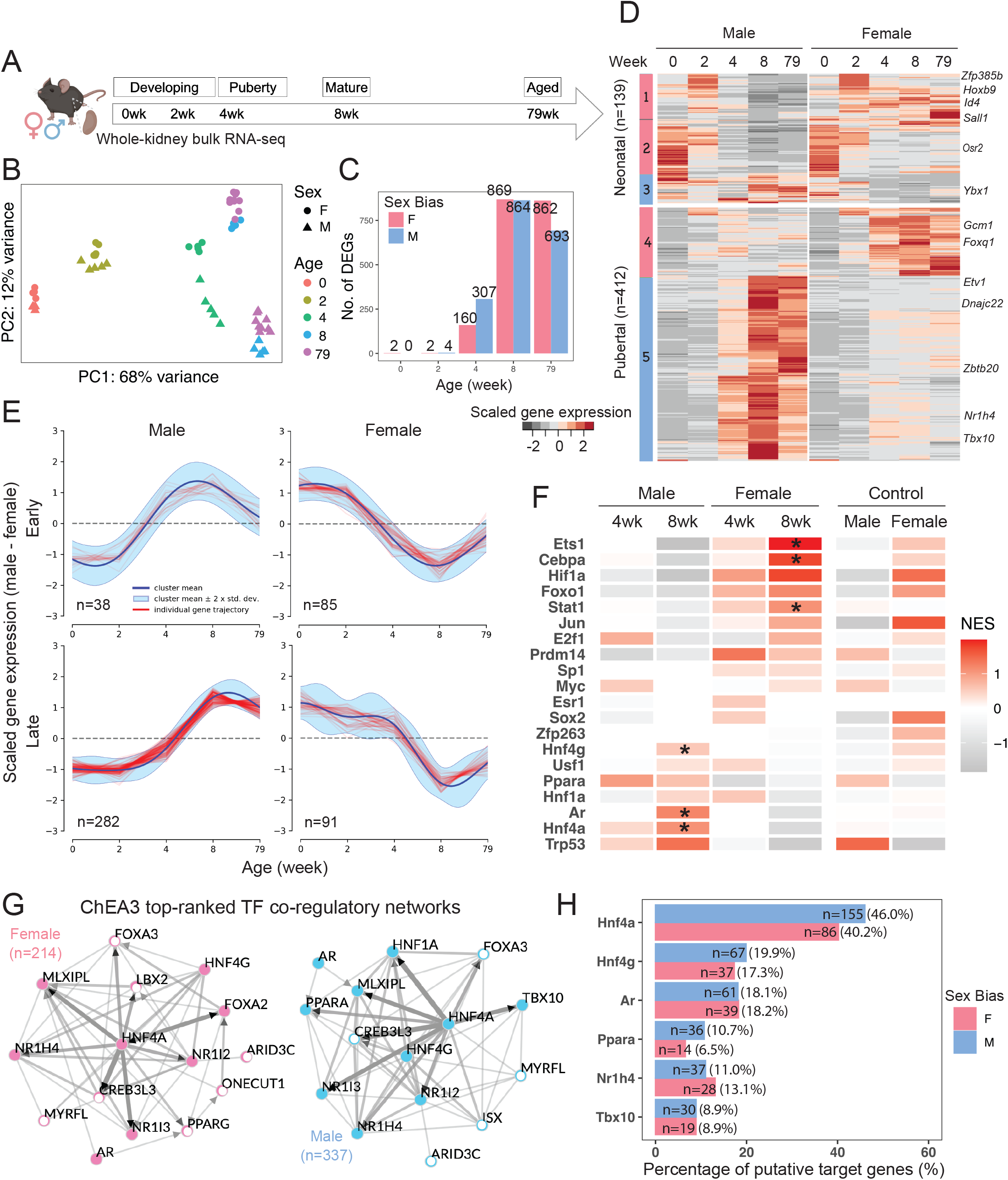
Male and female renal transcriptomes diverge at puberty. (A) Schematic summary of the experimental design in sampling renal transcriptome in male and female C57BL/6 mice. (B) Principal component analysis (PCA) reveals that the distribution of sample variations in gene expression are most influenced by age and sex. (C) Bar plot demonstrates the number of sex-biased genes identified at individual timepoints via differential expression analysis. (D) Heatmap shows the scaled average expression levels of the core sex-biased genes in male and female samples at individual timepoints. (E) Representative clusters of divergent gene expression dynamics analyzed by DPGP^57^. Red tracings represent genes, the navy line represents the mean divergent gene expression of the cluster, and the cyan margin shows the 95% confidence interval. (F) Tile map shows the predicted TF activities based on normalized gene expression in samples by DoRothEA^58, 59^, where high-confidence predictions are indicated by asterisks. (G) Network diagram of top 15 TFs that were predicted by ChEA3^60^ to regulate the core female and male programs. Edges indicate physical interaction supported by literature evidence, directed if supported by ChIP-seq data. Solid nodes indicate TFs that are expressed in the PT segments in our previous single-cell RNA-seq experiment^19^; open circles represent those that are not expressed. (H) Bar plot shows the percentage of putative targets among the core sex-biased genes that could be regulated by representative TFs, as predicted by ChEA3^60^. The number of putative targets is listed.

Comparing individual gene expression levels amongst the core gene set across all five timepoints revealed two broad categories of expression patterns (**Fig. 2D**): genes highly expressed at the earlier timepoints independent of sex (such as *Sall1* and *Bmp1*), representing developmental genes consistent with the continued differentiation and morphogenesis of the postnatal kidney (139 neonatal genes; 25%) and genes activated during puberty (4-8 weeks) many of which encode proteins participating in renal PT physiology (Table S1.2; 412 pubertal genes; 75%). Ninety percent of genes with M-biased expression (302 of the 337 M-biased genes) elevated expression during the 4-8 week period leading to sexual maturity, including transcriptional regulators such as *Nr1h4* and *Tbx10* (**Fig. 2D; Fig. S2G**). In contrast, only 51% of F-biased genes showed a similar trend in gene expression (**Fig. S2C**). In summary, for the male kidney, expression of most M-biased genes was induced, while about half of the F-biased genes were suppressed during puberty.

To examine the expression dynamics of the core sex-biased genes, we clustered the divergence of gene expression between male and female samples over time using DPGP^57^ (STAR Methods) to find two major patterns (**Fig. 2E**; Table S1.2): genes with expression diverging before 4 weeks (early) and those diverging on or after 4 weeks (late), which could signify the relative sequence of activation or inactivation. The anti-correlation of dynamic features among M-early and F-early genes (**Fig. 2E**) is suggestive of concomitant regulation during puberty, indicating that sex-biased gene expression in the male kidney could be driven by collaborating factors that activate the male program while simultaneously suppressing part of the female program.

To predict the upstream regulators for the divergence of male and female renal transcriptomes, we performed TF regulon analysis on high-confidence curated TFs using DoRothEA^58, 59^ (STAR Methods). Several TFs were predicted to specifically regulate the sex-biased program (**Fig. 2F**): AR (Ar), hepatocyte nuclear factor 1 alpha (Hnf1a), 4 alpha (Hnf4a) and gamma (Hnf4g) for M-biased program; CCAAT/enhancer-binding protein alpha (Cebpa), hypoxia induced factor 1 alpha (Hif1a), and JAK-STAT pathway for F-biased program. We noted slight enrichment for ERα (Esr1) activity at 4 weeks for activating female program, but not at 8 weeks (**Fig. 2F**).

To explore the role of less-known TFs, we ranked factors that potentially regulate the expression of the core sex-biased genes using ChEA3^60^. As expected, Hnf4a, as an important proximal tubular cell fate regulator during kidney development^61^, was centered as a hub that potentially regulated over 40% of both programs (**Fig. 2G-H**). We found that Hnf4a not only binds to and activate the expression of *Ar*, *Tbx10* and *Nr1h4* (ChIP-seq evidence^61^), but can also mediate the expression of M-biased genes (e.g., *Dnajc22*, *Ybx1*, and *Zbtb20)* and F-biased genes (e.g., *Gcm1* and *Foxq1)* (*in-silico* prediction^60^). AR could also regulate the expression of both programs (**Fig. 2H**), where there are substantial overlaps between putative targets of Hnf4a and AR (75 of 100, 75%; e.g., *Ppara*), consistent with previous findings^46^. In summary, these computational predictions suggest Hnf4a acts as a key upstream regulator for both male and female sex-programs in PT cells while AR plausibly regulates sex differences in the male kidney.

### The role of gonads, sex hormones, and sex hormone receptors in renal sex differences

Considering the emergence of renal sex differences at puberty and predicted involvement of AR regulation, we carried out a series of perturbation experiments (STAR Methods) to evaluate the role of AR, assaying responses through whole-kidney bulk RNA-seq (**Fig. 3A**). First, to understand the influence of the endogenous sex hormones, we performed prepubertal gonadectomy in mouse models (castration in males [CM] and ovariectomy in females [OF]) at 3 weeks and assayed kidney gene expression between the sexes at 8 weeks. Second, to investigate the effect of exogenous androgen, we injected testosterone subcutaneously into castrated males and ovariectomized females at 8 weeks, examining the kidney response 24 hours post-injection. Third, to study the role of sex hormone receptors directly on PT cells, we generated nephron-specific removal of Ar (Six2-Ar-KO) and Esr1 (Six2-Esr1-KO), then assayed kidneys at 8 weeks. Six2-CRE activity in nephron progenitor cells removes any potential sex hormone input from nephron progenitors and their nephron derivatives from the onset of embryonic kidney development^62^.

**Figure 3.**
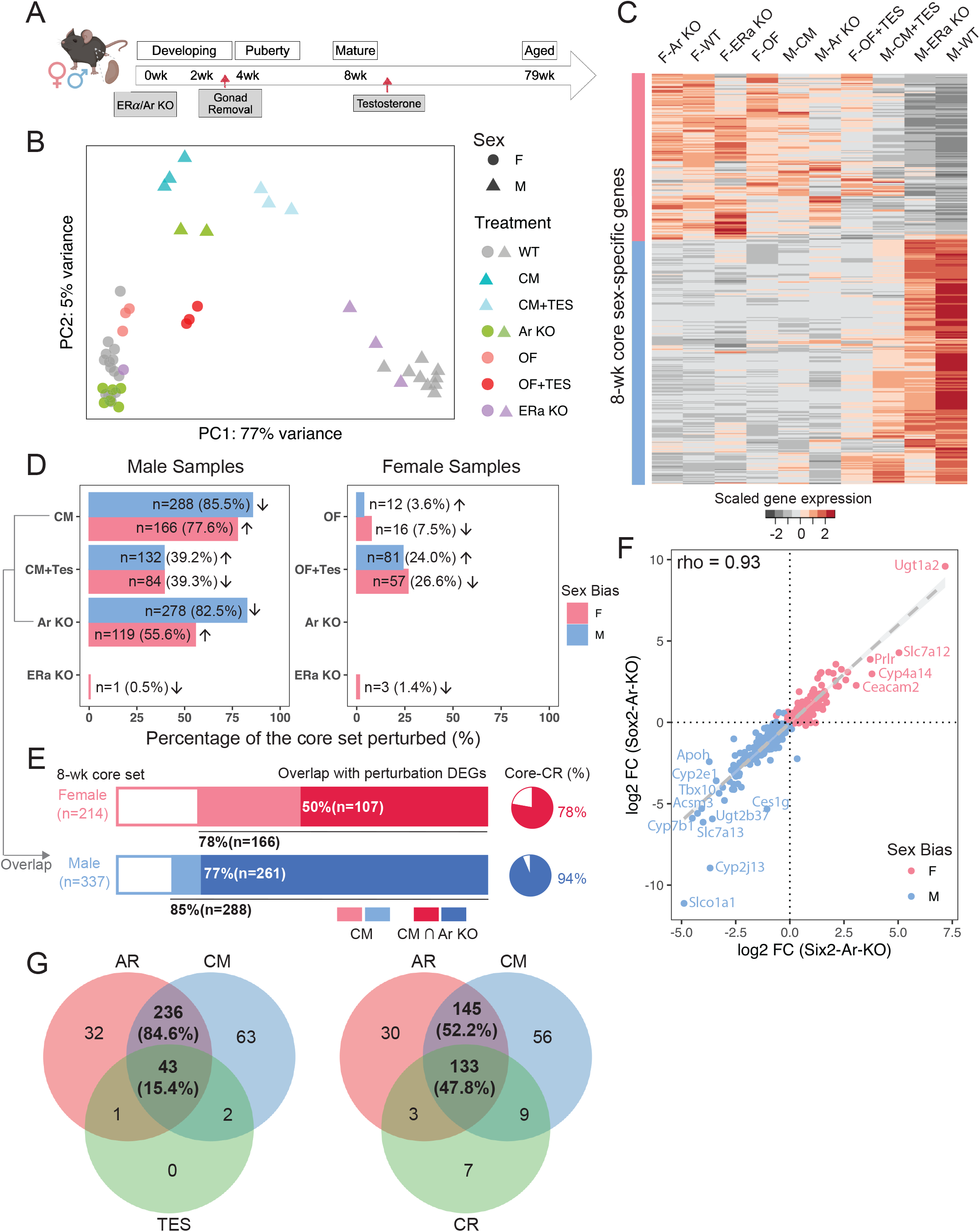
The role of gonads, sex hormones, and sex hormone receptors in renal sex differences. (A) The schematic summarizes the experimental design of perturbation treatments. Whole kidney bulk RNA-seq was performed between 8-12 weeks. (B) PCA plot demonstrates the relative renal transcriptional profile of mice undergoing various treatment regimens. Sample variations were evaluated based on the expression of the core sex-biased genes. CM: castration in males; OF: ovariectomy in females; TES: transient testosterone administration; WT: wild type; KO: knockout. (C) Heatmap shows the scaled average expression levels of the core set genes in male and female samples in individual treatments. (D) Percentage of the core sex-biased genes that were perturbed in individual treatments are shown for male and female samples in bar plots. The number of genes that are perturbed in each treatment and the corresponding percentages are listed. Arrows indicate the direction of perturbation in gene expression as compared to controls. (E) Stacked bars demonstrate the proportion of core sex-biased genes that are perturbed consistently in castration and nephron-specific AR knockout experiments in male samples. Pie charts show the percentage of AR-responsive genes that were perturbed in caloric restriction (CR) experiment^63^. (F) Scatter plot compares changes in the expression of core sex-biased genes in nephron-specific removal (Six2-Ar-KO) and systemic removal (Sox2-Ar-KO). (G) Venn diagrams showing overlap of perturbed core sex-biased genes in male kidneys among three groups: 1) AR: nephron specific removal of AR; 2) CM: castrated males; 3) CM+TES or CR: caloric restriction.

A comparative analysis of all the above against relevant control samples via PCA showed that castration and nephron-specific Ar removal had similarly strong effects: partially feminizing the male kidney (**Fig. 3B-C**). After castration, 85.5% of the core M-biased genes (288 of 337) were down-regulated and 77.6% of core F-biased genes (166 of 214) were up-regulated (**Fig. 3D; Fig. S3A**). On nephron-specific Ar removal, a comparable number of sex-biased genes showed a significant change in expression: 82.5% of core M-biased genes (278 of 337) were down-regulated while 55.6% of core F-biased genes (119 of 214) were up-regulated. Most (93.9%; 261 of 278) of the Ar-dependent M-biased core gene set were also down-regulated in the castration experiment (**Fig. 3E**). Transient administration of testosterone partially restored the male phenotype following castration and activated a male-like program in ovariectomized females (**Fig. 3B-D**; Table S2.1-4). In contrast, nephron removal of either *Ar* or *Esr1* had little effect on gene expression in the female kidney in line with ovary removal and computational predictions (**Fig. 2F**; **Fig. 3B-D**; Table S2.5-8).

Comparing male kidneys following systemic whole-body AR removal through CRE-mediated recombination at embryonic implantation (Sox2-Ar-KO; Table S2.9) with nephron-specific removal (Six2-Ar-KO) showed a high concordance in the male kidney gene expression datasets (Spearman correlation rho=0.93; **Fig. 3F**), comparable to male-female comparisons (**Fig. S3B-C**). Thus, systemic removal of *Ar* was equivalent to local removal of *Ar* in the nephron consistent with direct action of AR in proximal tubule cells. A list of AR-responsive genes is presented in Table S2.10. M-biased genes such as *Slco1a1*, *Cyp2j13, Cyp7b1, Slc7a13, Acsm3*, and *Tbx10,* were significantly down-regulated in both nephron-specific and systemic removal of AR; F-biased genes *Ugt1a2, Scl7a12, Prlr*, and *Cyp4a14* were most up-regulated (**Fig. 3F**). Interestingly, *Slco1a1* and *Cyp2j13* showed a significantly larger fold change in systemic versus nephron-specific AR removal, suggesting that extra-renal action of AR (e.g., via the hypothalamic-pituitary axis) also regulates their expression. In summary, these findings provide compelling evidence that the primary determinant of sexually dimorphic gene expression is the presence (male) or absence (female) of testosterone-mediated regulation of AR activity in PT cells of the mouse kidney.

In addition, by comparing against a published kidney RNA-seq dataset on caloric restriction^63^ (CR), we found that 47.8% of the core sex-biased genes that were perturbed by both castration and AR removal were altered in male mice after a regimen of 3-month reduced food intake (**Fig. 3G**). Moreover, 78% of CR-up-regulated genes are AR-responsive (F-biased) and 94% of CR-down-regulated genes are AR-responsive (M-biased) (**Fig. 3E**), indicating that CR modifies the sex profile of the kidney.

### Single-nuclear multiomic profiling of AR function in the mammalian kidney

To obtain a more detailed insight into AR regulation in the nephron, we applied 10X multiomic single-nuclear RNA-and ATAC-seq profiling to examine chromatin regulation and aligned gene activity in WT and Six2-Ar-KO kidneys at 8-10 weeks of age (**Fig. 4A**). Stringent quality control steps and depth normalization approaches were applied to minimize technical and batch effects (STAR Methods). Data from individual samples were integrated and nuclei were clustered based on both RNA and ATAC modalities (STAR Methods). Clusters were manually annotated based on established cell-type markers (**Fig. S4A-B**).

**Figure 4.**
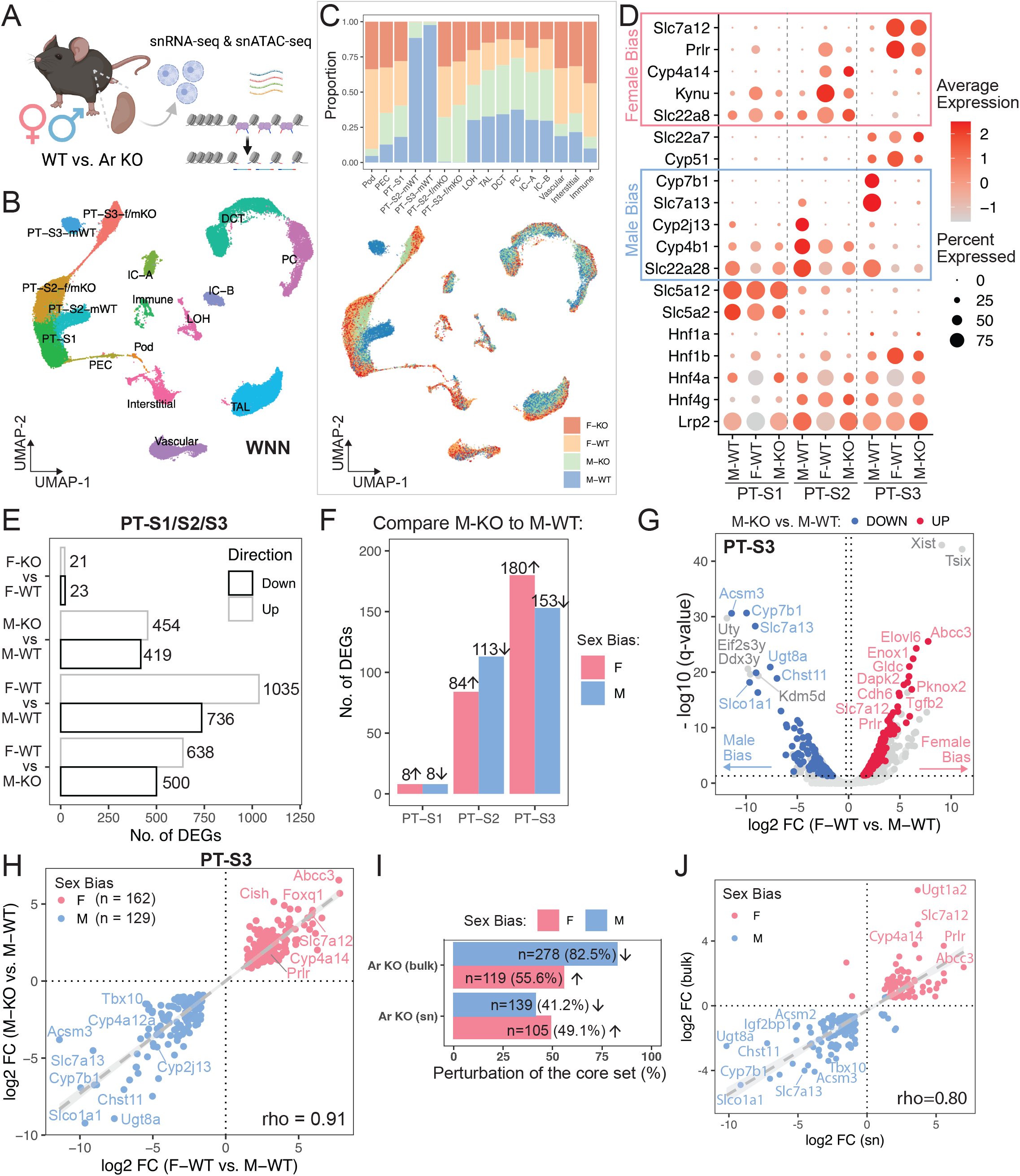
Single-nuclear multiomic profiling of AR function in the mammalian kidney. (A) Schematic summary of the single-nuclear multiomic experiment. (B) UMAP plot indicates the divergent features between male and female PT cells while the other cell populations co-cluster regardless of sex. Nuclei were clustered based on RNA and ATAC modalities using weighted nearest neighbor (WNN) graph. (C) Distribution of sex and genotype among all cell populations shown in (B). Top: stacked bar plot shows composition in each cluster; bottom: nuclei in the UMAP plot (B) colored by different sex-genotype combinations. (D) Dot plot demonstrates the expression pattern of top marker genes for individual PT segments. Known sex-biased genes are indicated. (E) Bar plot shows the total number of differentially expressed genes identified using the multiomic RNA data by four pairwise comparisons within PT segments. (F) Bar plot lists segment-wise number of single-nuclear sex-biased genes that were perturbed upon AR removal. (G) Volcano plot shows single-nuclear sex-biased genes identified in PT-S3 segment, where genes that are perturbed by AR removal in the male kidney are highlighted. (H) Scatter plot contrasts the effect of nephron-specific AR removal in male to the observed sex biases. (I) Bar plot compares the percentage of the core sex-biased genes that were perturbed by nephron-specific AR removal, between bulk and single-nuclear RNA-seq. (J) Scatter plot shows the impact of nephron-specific AR removal on common sex-biased gene, using bulk or single-nuclear RNA-seq data.

As expected, integrated RNA/ATAC data suggest molecular differences between sexes and genotypes predominantly manifest among PT cell clusters, where previous single cell RNA-seq studies^19^ have demonstrated sexually dimorphic gene activity predominantly maps to PT segments 2 and 3 (PT-S2 and PT-S3; **Fig. 4B**). In the multiomic data, PT segment 1 (PT-S1) cluster comprised S1 cells from all genotypes and sexes, indicative of a low level of AR-dependent variability between male and female sexes (**Fig. 4B**), albeit with a slight segregation of male and female samples (**Fig. 4C**), which may be due to residual sequencing imbalances not accounted for during batch correction. In contrast, WT nuclei from PT-S2 and PT-S3 clustered separately comparing male and female kidney samples (**Fig. 4B, C**). Further, the vast majority of male Six2-Ar-KO nuclei clustered with female WT and Six2-Ar-KO nuclei (**Fig. 4B-C**). Analysis of the expression of top male-and female-specific markers distinguishing individual PT segments indicated male PT segments resemble their female counterparts following AR removal (**Fig. 4D**). These data indicate that each PT segment adopts a distinct segmental identity, with a segment-specific female ground-state that is masculinized by the direct action of Ar in response to androgens in the male kidney.

For a systematic evaluation of transcriptomic profiles between these sex-genotype combinations, we performed differential expression analysis within individual PT segments (STAR Methods). Using the single-nuclear RNA data, we identified a total of 1,035 F-biased and 736 M-biased genes (**Fig. 4E**; Table S3.1; representing the “single-nuclear sex-biased genes”), where higher sequencing depth in female samples possibly inflated the number of F-biased genes. Of note, PT-S3 showed the highest number of sex-biased genes (**Fig. S4C**). Upon AR removal in male nephrons, 220 F-biased genes were up-regulated among PT segments while 211 M-biased genes were down-regulated (Table S3.2; see **Fig. 4F** for segment-specific quantification), which comprise a set of single-nuclear sex-biased genes with large effect sizes (**Fig. 4G**; **Fig. S4F**); but the expression of sex-chromosome genes was independent of AR. These AR-responsive sex-biased genes showed high concordance of relative gene expression in female WT and male KO PTs when compared to male WT (Spearman correlation rho=0.91; **Fig. 4H**; **Fig. S4G-H**). Among AR-responsive sex-biased genes are *Slco1a1* and *Abcc3*, showing the largest differences in comparing male and female WT PT-S3, respectively, as well as the largest changes in expression before and after AR removal in the male PT-S3 (**Fig. 4H**). The small number of gene expression changes observed on AR removal from female nephrons (21 genes up-regulated and 23 down-regulated) likely represent background effects in the data rather than *bone-fide* regulation (**Fig. 4E**; Table S3.3). When we compared changes in expression before and after AR removal between bulk and single-nucleus RNA-seq experiments, we found that 41-49% of the core sex-biased genes identified in bulk data (**Fig. 1B**) were identified in the single-nuclear data (**Fig. 4I**), which also show a high concordance in fold change (Spearman correlation rho=0.80; **Fig. 4J**).

### snATAC identified AR response elements near sex-biased genes

To further delineate the molecular mechanism of AR-directed regulation of dimorphic gene expression, we examined the chromatin landscape within each kidney cell type using the single-nuclear ATAC data, with a focus on PT cells (**Fig. 5A**). Removal of *Ar* resulted in a pronounced co-clustering of *Ar* mutant male PT cells with wild-type and *Ar* mutant female PT cells suggesting AR plays a major role in sex-biased regulation of chromatin accessibility in PT (**Fig. 5A**). Global comparison of chromatin states between PT populations revealed that segment-specific differences are larger than sex-dependent differences within each segment (**Fig. S5A**). We applied differential accessibility analysis between male and female WT nuclei to identify potential functional response elements that are proximal or distal to the AR-responsive genes (Table S4) (STAR Methods). Akin to sex-biased gene expression (**Fig. S4C**), PT-S3 showed the highest number of differentially accessible regions (sex-biased DARs; **Fig. 5B, S5B**): 7,987 F-biased peaks and 11,972 M-biased peaks, with 1,160 F-biased peaks and 1,087 M-biased peaks within 100KB of the transcriptional start site (TSS) of genes with sex-biased expression (Table S3.4). Pairwise comparison of DARs across sex and genotype detected fewer DARs in M-KO vs M-WT PTs than M-WT vs F-WT DARs, leaving open the possibility for AR-independent regulation of chromatin in male PT cells (**Fig. 5B**).

**Figure 5.**
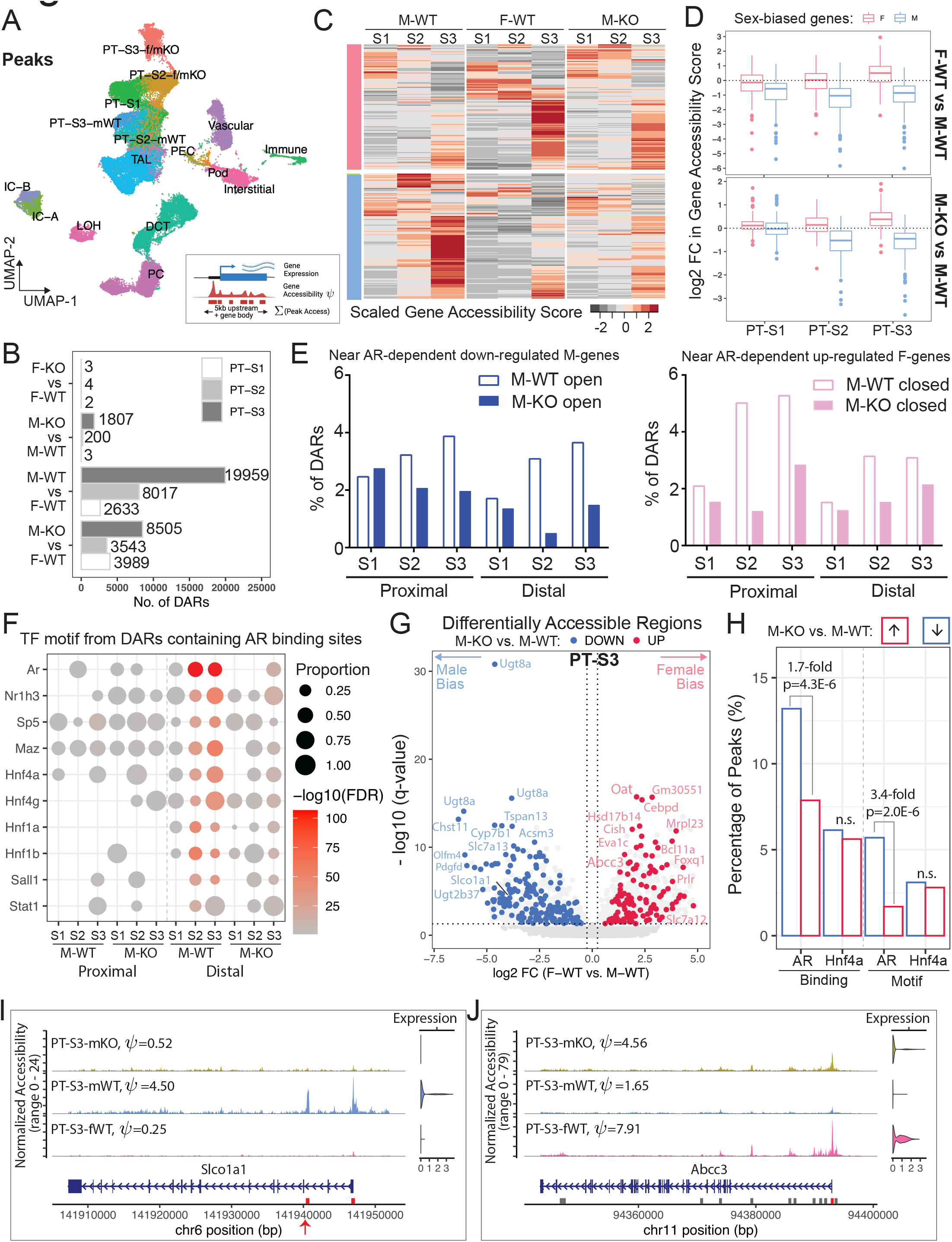
AR response elements are located near sex-biased genes. (A) UMAP plot shows clustering outcome using peaks called from the single-nuclear ATAC data. PT-S3-f/mKO: co-clustering of PT-S3 cells from F-WT, F-KO, and M-KO; PT-S2-f/mKO: co-clustering of PT-S2 cells from F-WT, F-KO, and M-KO; mWT: M-WT. (B) Bar plot shows the number of sex-biased Differentially Accessible Regions (DARs) in PT segments identified for each pair-wise comparison (absolute Log2FC > 0.25, adjusted p-value < 0.05). (C) Schematic summary of the computation of gene accessibility score *ψ* and heatmap shows the scaled gene accessibility score *ψ* for AR-responsive genes in M-WT, F-WT, and M-KO PT segments. (D) Box plots demonstrate fold change in gene accessibility score *ψ* of AR-responsive genes within individual PT segments. (E) Histograms showing the percentage of the proximal and distal DARs from M-WT and M-KO compared to F-WT that were nearby AR dependent down-regulated M-biased (blue) and up-regulated F-biased (pink) genes. (F) Dot plot summarizes TF motif enrichment in the open DARs containing AR binding sites based on the published CHIP-seq dataset^46^. in M-WT and M-KO compared to F-WT PT segments. (G) Volcano plot shows DARs within 100KB of sex-biased genes in PT-S3. 11,972 peaks were differentially open in male (left) and 7,987 peaks were differentially open in female (right). We colored F-biased peaks that are preferentially open in M-KO in red, and M-biased peaks that are preferentially closed in M-KO in blue. Each dot represents a 500- bp region, where the nearest gene is annotated. (H) Bar plot shows the prevalence of TF binding and motif among PT-S3 sex-biased DARs that were altered by AR removal. TF binding was based on ChIP-seq data in the mouse kidney^46^. TF motif PWMs were retrieved from the Jasper database^108^ for AR (MA0007.3) and Hnf4a (MA0114.3). (I-J) Coverage plots of two representative sex-biased genes, *Slco1a1* (M-biased; I) and *Abcc3* (F-biased; J). All peaks called in the region are shown in gray boxes, where DARs are highlighted in red. Peaks with potential AR binding site are indicated by red arrows.

To examine AR dependent sex-specific chromatin differences, we compared M-biased open and closed (the latter equivalent to F-biased open) DARs between M-WT and M-KO using F-WT as a common standard (**Fig. S5B**, Table S4). Of the M-biased open peak set from S2 (11079 peaks) and S3 (15307 peaks), 83.1% of the S2 peaks and 66.7% of the S3 peaks showed a loss of differential accessibility comparing M-KO to F-WT in line with direct AR regulation of chromatin accessibility (**Fig. S5B**). Examining the F-biased open peaks, approximately half gained female-like accessibility on AR removal in the male kidney (**Fig. S5B**).

For the AR-responsive sex-biased genes identified from the multiomic data (**Fig. 4F-H, Fig. S4F-H**), we evaluated their chromatin state using gene accessibility score *ψ* (**Fig. 5C**), a metric that quantifies the openness of a genomic region using a weighted sum of peaks within the gene body and promoter region (up to 5KB upstream of its TSS; STAR Methods). As expected, F-biased genes are preferentially open in female PTs, while M-biased genes are more open in male PTs (**Fig. 5C-5D**). Following AR removal from male nephrons, 98% of M-biased genes show decreased accessibility, especially in PT-S2 (**Fig. 5C-5D**; Table S3.5), while 97% of F-biased genes showed a more open chromatin profile, mostly prominent in PT-S3 (**Fig. 5D**; Table S3.5). In male PT-S1, little effect was observed on either gene expression (**Fig. S4D**) or chromatin status (**Fig. 5D**) upon AR removal. Consistently, the most striking reduction in the DARs near AR dependent down-regulated genes in M-KO were found in the male-biased open distal regions in PT-S2, and in those near AR dependent up-regulated female genes in male-closed proximal regions (within 1KB upstream of TSS) in PT-S2 (**Fig. 5E**). For genes with a F-bias, *Ar* removal in the male kidney resulted in an increase in open chromatin in proximal and distal regions in both PT-S2 and PT-S3 (**Fig. 5E**), consistent with the activation of a F-biased gene set (**Fig. 4G**).

AR binding to chromatin associated with kidney target genes has been identified through ChIP-seq following acute testosterone administration injection^46^. To examine the AR dependent transcription factor binding in the male open DARs regions compared to F-WT, we performed motif enrichment analysis on DARs associated with Ar binding in the CHIP-seq experiments. This analysis showed a strong enrichment for predicted Ar motifs in distal regions associated with cis-regulatory elements, as well as motifs for several other transcriptional regulators, notably Hnf1a/1b and Hnf4a/4g, which are broad regulators of proximal tubule programs (**Fig. 5F**). Thus, PT specific actions of Ar are likely to be directed by a broader PT gene regulatory program.

We also performed differential accessibility analysis between M-WT and M-KO nuclei (Table S3.6). Among the sex-biased DARs identified (**Fig. 5B**), 167 F-biased peaks became more open upon AR removal in male PT-S3, while 211 M-biased peaks became more closed (**Fig. 5G**; AR-responsive DARs). Notably, many of the AR-responsive DARs were near genes with the most marked sex-biased expression (see also **Fig. 4G-H**). When AR-responsive PT-S3 DARs were compared with the AR ChIP-seq data^46^, we observed a 1.7-fold increase in AR binding among M-biased peaks than F-biased peaks (two-proportion z-test, *P*-value=4.3E-6), whereas Hnf4a binding did not show a significant difference (**Fig. 5H**). When compared to F-biased peaks, AR-responsive M-biased peaks showed an enrichment for AR motif (two-proportion z-test, *P*-value=2.0E-6), but not for the Hnf4a motif (**Fig. 5H**). PT-S2 DARs behaved similarly, but none of the sex-biased DARs in PT-S1 were perturbed on AR removal (**Fig. S5D-E**). Together, these data are consistent with direct AR regulation of cis-regulatory modules driving expression of male-enriched gene expression in the S2 and S3 segments of the PT.

To identify additional TFs that mediate the sex-biased transcription program, we performed TF motif enrichment analysis on sex-biased DARs within 100KB of the TSS of the AR-responsive gene set (**Fig. S5C**, Table S3.7). In addition to Ar, M-biased DARs predicted motif enrichment in PT-S3 for Rfx3, a key factor in cilium biogenesis^64^. Motif recovery further highlighted the likely interface of Ar action with general PT regulatory programs mediated by Hnf1a/b and Hnf4a/g which were enriched in in both M-and F-biased DARs (**Fig. S5C**; Table S3.7). F-biased DARs showed a strong motif enrichment for Stat5a/b and Bcl6 which lie downstream of prolactin and growth hormone signaling^65, 66^ suggesting alternative pathways of F-biased regulation (see discussion).

Integrating AR ChIP-seq^46^ and our multiomic data, we were able to uncover putative AR response elements near sex-biased genes. For example, there were 5 M-biased DARs (2 proximal and 3 distal) annotated in the genomic region of *Slco1a1* in PT-S3 (Table S3.4), and 3 were preferentially closed in male PT-S3 when AR was removed (Table S3.6), including the intronic peak with AR binding site (**Fig. 5I**), a phenomenon that was also found in PT-S2 (**Fig. S5F**). Note that out of the 6 M-biased DARs (5 intronic and 1 in the promoter) with AR binding annotated in the genomic region of *Cyp2j13*, the promoter region peak was identified as the most significantly down-regulated post AR removal (FDR<3.4E-5) (**Fig. S5G;** Table S3.4, S3.6). In the genomic region of *Abcc3*, we detected 11 F-biased DARs (6 within the gene body and 5 distal) in PT-S3 (Table S3.4); one peak at the promoter was preferentially open in male PT-S3 without AR (**Fig. 5J**), and another peak 53KB upstream in between Abcc3 and Cacna1g was bound by both AR and Hnf4a (Table S3.4, S3.6), suggesting multi-factor regulation of sex-biased gene expression. Collectively, examination of AR responsive peaks near candidate genes with AR binding information provided strong evidence for direct AR action mediating chromatin changes near M-biased genes in PT-S2 and S3, but not F-biased genes.

### RNAscope validates dimorphic gene expression in proximal tubule

To visualize gene expression directly in PT segments, we combined uniquely labeled RNAscope probes and performed *in situ* hybridization to adult male and female kidney sections (**Fig. 6**). M-biased gene *Cyp2j13* (a member of Cytochromes P450 family, metabolizing arachidonic acid into epoxyeicosatrienoic acids for vasodilation and other functions^67^) exhibited AR-dependent expression in PT-S2, colocalized with S2 marker *Cyp2e1* (**Fig. 6A**). *Slco1a1* is the top candidate for AR-responsive M-biased gene (**Fig. 4H**), which encodes solute carrier organic anion transporter polypeptide 1 (OATP1) important for the uptake of steroid conjugates and prostaglandin E2 into the cell^68, 69^. As predicted, *Slco1a1* mRNA was highly male-specific among PT-S2, and entirely absent upon AR removal from male nephrons (**Fig. 6B**). M-biased *Atp11a* encodes phospholipid-transporting ATPase IH, an integral membrane P4-ATPase that function as flippases at the plasma membrane to translocate phospholipid from the outer to the inner leaflet^70^. *Atp11a* expression was M-biased in PT-S2 and PT-S3; total mRNA in individual PT cells recapitulated single-cell measurement (**Fig. 6C**). Interestingly, *Atp11a* mRNA was abundant in the cytoplasm of male PT-S2 and PT-S3 but was concentrated in the nuclei among female PTs as well as among male PT cells without AR (**Fig. 6C**). Phospholipid-transporting ATPase IH is reported to be actively translated only in male PTs, where phospholipid asymmetry across the cell membrane regulates solute transport and membrane protein function^71, 72^.

**Figure 6.**
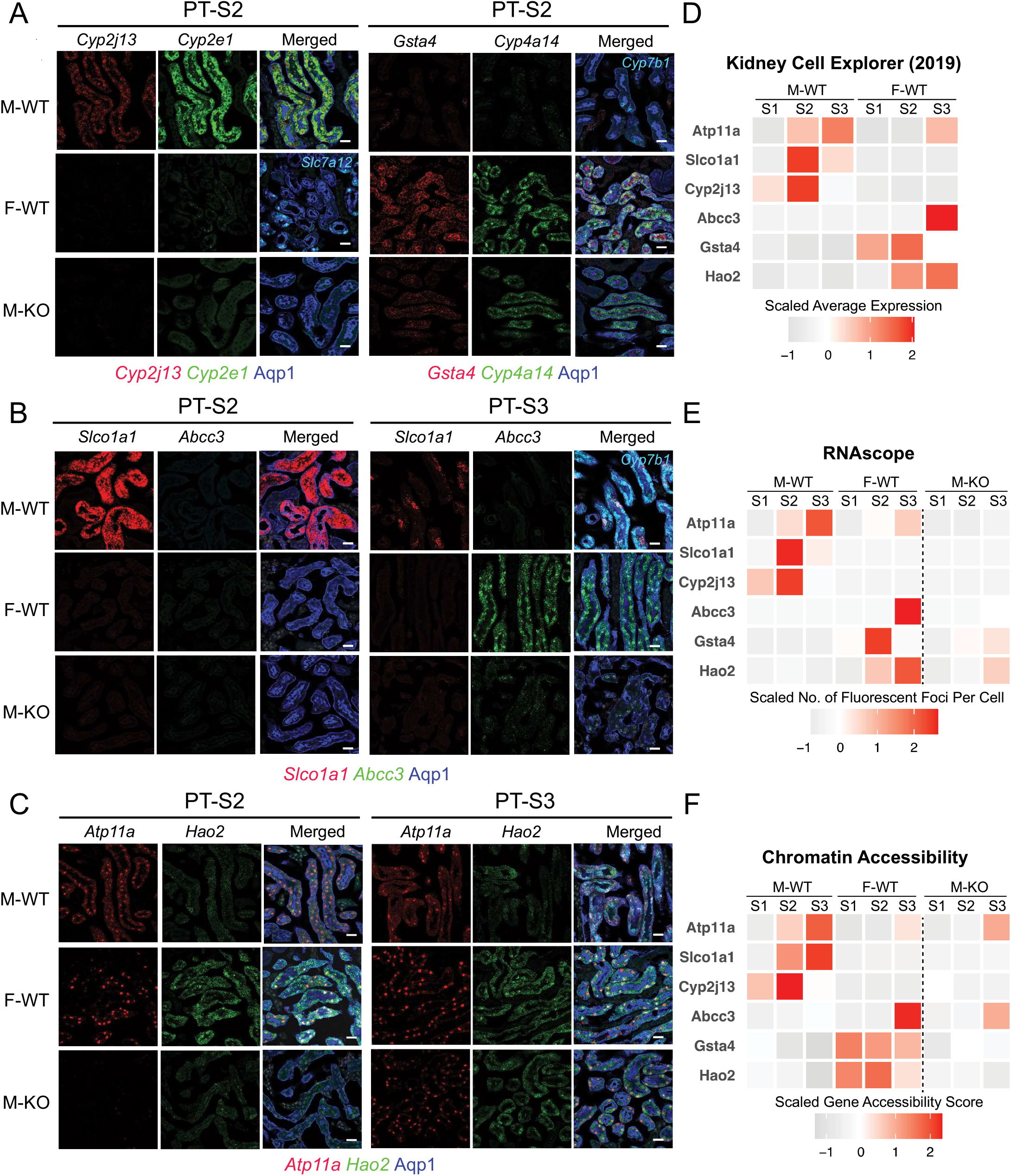
Fluorescence RNA in situ hybridization by RNAscope validates dimorphic gene expression in proximal tubule. (A-C) RNAscope assay directly visualized the expression levels of sex-biased genes in M-WT, F-WT, and M-KO PT-S2 & S3 (scale bars = 20 μm): co-stained with an antibody against Aqp1 (blue) demarcating the PT. (A) Left: *Cyp2j13* (red) and *Cyp2e1* (green, male PTS2 marker) co-stained with *Slc7a12* (Cyan, female PTS3 marker); right: *Gsta4* (red) and *Cyp4a14* (green, female PTS2 marker) co-stained with *Cyp7b1* (Cyan, male PTS3 marker). (B) *Slco1a1* (red) and *Abcc3* (green), co-stained with *Cyp7b1* (Cyan, male PTS3 marker) (C) *Atp11a* (red) and *Hao2* (green). (D-F) Tile maps show the expression and chromatin profile of top sex-biased genes in M-WT and F-WT PT segments: (D) data from previous scRNA-seq experiment^19^; (E) *in-situ* expression of top sex-biased genes measured by RNAscope; and (F) the estimated chromatin accessibility.

The F-biased gene *Gsta4* (Glutathione S-transferase alpha 4), which is known to protect against oxidative injury and renal fibrosis^73^), was highly differentially expressed in female PT-S2 and induced in the male PT-S2 segment on AR removal (**Fig. 6A**). The top F-biased gene *Abcc3 (*or *Mrp3)* encodes a member of the superfamily of ATP-binding cassette (ABC) transporters, essential for the efflux of organic anions, including steroid conjugates, glutathione conjugates and prostaglandin J2^74^. *Abcc3* showed female-specific expression in PT-S3 comparing male and female kidneys, and a striking, though partial, up-regulation upon AR removal from male nephrons (**Fig. 6B**). In addition, the F-biased gene *Hao2* encodes peroxisomal hydroxy acid oxidase 2, which was shown to eliminate lipid accumulation and inhibit progression of clear cell renal cell carcinoma^75, 76^. *Hao2* was highly expressed in female PT-S2 and PT-S3, but only low levels of expression were detected in homologous male PT segments (**Fig. 6C**); a marked increase in PT-S3 *Hao2* expression was observed on AR removal from the male nephrons (**Fig. 6C**). Together, RNAscope experiment validated the expression pattern of candidate AR-responsive sex-biased genes in PT segments, as predicted by sequencing results (**Fig. 6D-E**). Further, we found that the chromatin accessibility profile of these candidate genes was also largely in line with the expression pattern (**Fig. 6F**).

### Distinct and shared processes of dimorphic gene expression in the kidney and liver

In contrast to the kidney, the liver has been shown to be regulated by sex-dependent dynamics of growth hormone stimulation, where growth hormone release by the hypothalamus-pituitary axis is under direct androgen and estrogen control^77, 78^. However, to our knowledge, the effects of direct AR removal from hepatocytes on sexually dimorphic expression in the liver have not been addressed. To compare liver and kidney mechanisms, we initially identified a total of 1,682 genes with sex-biased gene expression in the C57BL/6 mouse liver at 8-12 weeks through bulk RNA-seq (**Fig. 7A**; Table S5.1), a comparable number to the 1,733 sex-biased genes identified in the kidney at 8 weeks (**Fig. 1B**). This list of hepatic sex-biased genes covers 43-55% of those identified in two previous studies^79, 80^ analyzing bulk RNA-seq of 8- (GSE174535) and 16-week (GSE112947) liver samples (**Fig. 7A**). The concordance of expression profile between datasets was high (Spearman correlation rho=0.92-0.93; **Fig. S7B**). Comparing sex-biased genes in the liver and kidney, we identified 102 shared M-biased genes (15%) and 143 shared F-biased genes (15%) (**Fig. 7B**), a significant conservation of sex differences between the liver and kidney (Chi-square test; *P*-value = 2.2E-16), when compared to genes shared across sexes in the two organs (5%).

**Figure 7.**
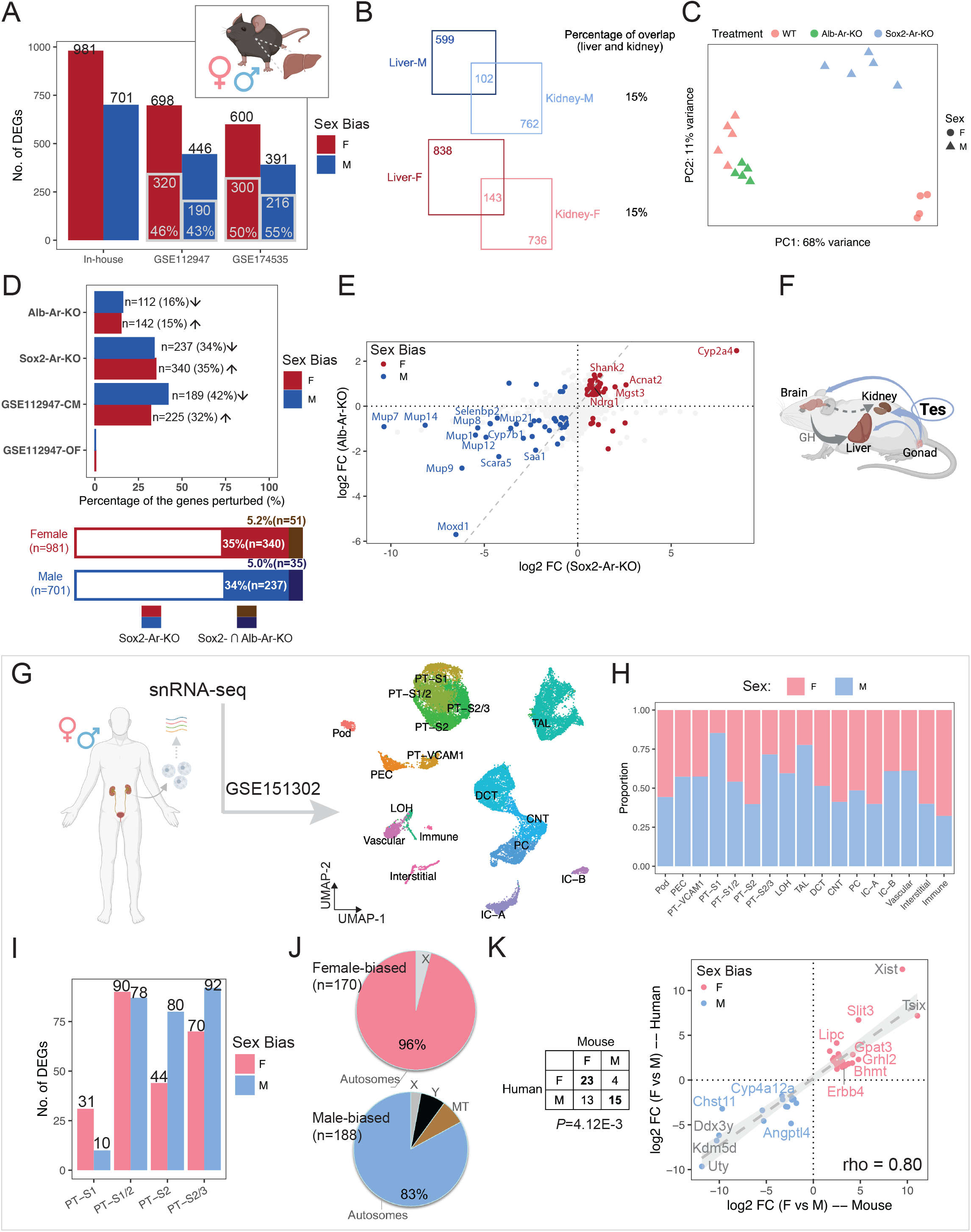
Distinct and shared processes of dimorphic gene expression between organs and species. (A) Bar plot shows the number of liver sex-biased genes identified in the current study (in-house) and those reported in the literature. Gray-contoured bars indicate the number of genes overlapping with the in-house list. (B) Venn diagrams show the number of sex-biased genes that are shared in the kidney and liver. (C) PCA plot demonstrates the impact of hepatocyte-specific (Alb-Ar-KO) and systemic AR removal (Sox2-Ar-KO), as compared to WT samples. (D) The percentage of in-house liver sex-biased genes that were perturbed in individual treatments is shown in the bar plot (top) and stacked bar plot (bottom). Arrows indicate the direction of perturbation in gene expression when compared to controls. (E) Scatter plots compare the changes in expression of in-house liver sex-biased genes between systemic and hepatocyte-specific AR removal. The dashed gray diagonal line marks equal impact. (F) A schematic summary of how testosterone influences the sexual dimorphism in the kidney and liver. (G) UMAP plot shows clustering of human renal snRNA-seq data (GSE151302). (H) Composition of male and female cells in each cluster in (B). (I) Bar plot shows the number of sex-biased genes among each PT cluster in (B). (J) Pie charts demonstrates the percentage of autosomal versus X/Y-linked genes among all the sex-biased genes identified in (D). (K) Comparison of sex-biased gene expression in human and mouse kidney reveals conserved sexual dimorphism. The table lists the number of orthologs that show sex biases in gene expression; the scatter plot shows the differences in expression of common sex-biased genes in human and mouse PT segments.

To investigate the role of AR in the mouse liver, we compared hepatocyte-specific removal of AR with an albumen CRE strain^81^ (Alb-Ar-KO) to systemic removal (Sox2-Ar-KO). As expected, PCA analysis across perturbations showed systemic AR removal had a more dramatic effect than hepatocyte-specific removal (**Fig. 7C-D**). Systemic AR removal resulted in the down-regulation of 34% of genes with M-biased expression and up-regulation of 35% of those with F-biased gene expression (**Fig. 7D**; Table S5.2), consistent with previously reported castration experiments^79^ (**Fig. S7C-D**). In contrast, 15-16% of sex-biased genes were perturbed on hepatocyte-specific AR removal, in directions consistent with male/female biases (**Fig. 7D**; Table S5.3). 5% of genes with liver sex biases in expression were shared between the systemic and hepatocyte-specific AR removal (**Fig. 7D**). Further, the magnitude of expression changes was greater for systemic AR removal (**Fig. 7E**), with a particularly marked alteration in the expression of genes encoding major urinary proteins (*Mup1/7/8/9/12/14/21*) and cytochrome P450 member (*Cyp2a4*). We did not detect a high concordance between systemic and hepatocyte-specific AR removal (Spearman correlation, rho=0.39; **Fig. 7E**), indicating the indirect action of AR controlling sex-specific gene expression in hepatocytes: a moderate perturbation was observed following AR removal in hepatocytes to a small number of genes sharing sex-biased expression in the liver and kidney. Only a handful of genes were predicted to share direct AR control of their expression in these two organs, including *Cyp7b1, Selenbp2, Cyp4a12a and Oat* (Table S5.4).

### Conserved renal sex differences in the human and mouse

Several reports have documented human kidney-associated differences in gene expression between the sexes^82–84^. To determine whether conserved mechanisms extend from the mouse to the human kidney, we re-analyzed bulk RNA-seq data on kidney biopsies from adult male and female donors of diverse demographic background (GTEx v8^83^; STAR Methods). These studies showed only a modest number of genes differentially expressed between the sexes in human, 1 F-biased gene (*KDM5C*, X-linked) and 73 M-biased genes (2 X-linked, 63 Y-linked, and 8 autosomal). The three conserved sex-biased genes comparing mouse (both the core and full set; **Fig. 1B**; Table S6) and human data are all encoded by the Y chromosome: *UTY, DDX3Y*, and *KDM5D*.

Examining recent snRNA-seq dataset for the human kidney (2 female and 3 male donors; GSE151302^85^; **Fig. 7G**; STAR Methods) showed co-clustering of expression data for male and female PT segments with no apparent sex bias in cluster composition (**Fig. 7H**). Differential gene expression analysis uncovered a total of 170 F-and 188 M-biased genes, over 80% of which are autosomal (**Fig. 7I-J**; Table S6). Through ortholog matching, we identified 23 F-biased genes and 15 M-biased genes with conserved expression between the human and mouse kidney (**Fig. 7K**; Fisher’s exact test, *P*-value=4.12E-3), including predicted AR-responsive genes in the murine kidney (*Chst11* and *Bhmt*; Table S2 & S3) and murine kidney & liver (*Cyp4a12a*; Table S5.4). Though there are several caveats with these human studies (see discussion), these findings suggest a limited conservation in sex-biased expression and AR-mediated in-organ regulation between the mouse and human kidney.

## Discussion

In this study, we used time-course bulk RNA-seq and single-nuclear multiomic data to investigate the regulatory mechanism of renal sexual dimorphic gene expression in mice. Sexually dimorphic gene expression in PT cells is established under gonadal control between 4 to 8 weeks postpartum primarily through androgen signaling; ovary removal and Esr1 deletion had little effect. Several lines of evidence support a direct regulatory action of Ar binding to chromatin within cis-regulatory regions of genes showing M-biased expression as a major driver of sexually dimorphic gene expression in the mouse kidney. Critically, androgen receptor activity in PT cells is required for establishing a normal male program of gene expression. The requirement correlates with androgen responsiveness of M-biased genes, and Ar motif enrichment and Chip-seq binding studies, that point to Ar engagement within distal regulatory regions of genes with M-biased expression. Co-recovery of motifs for general regulators of PT identity and cell function (Hnf1a, Hnf1b, Hnf4a, Hnf4g) suggests Ar acts is conjunction with broader PT regulatory mechanisms.

While there is strong evidence for a direct activating role for Ar in controlling M-biased genes, the mechanisms regulating F-biased gene expression are less clear. Loss of Ar in PT cells results is a substantial ectopic activity of F-biased genes indicating that suppression of the female program is dependent on direct Ar activity in PT cells. However, we did not observe a strong enrichment of Ar motifs in distal regulatory regions around the F-biased gene set suggesting an indirect regulatory role; for example, transcriptional activation of a gene encoding a repressor of the female program. The F-biased program is also associated with motif predictions from DARs for Stat5a, Stat5b and Bcl6, a negative regulator of Stat action^41^. Interestingly, both male and female patterns of sexually dimorphic gene expression in the mouse liver are controlled through growth hormone signaling to hepatocytes^44^, though our analysis of hepatocyte removal of *Ar* suggests a minor role for direct Ar action (see below). These findings raise the possibility of direct growth hormone control of the F-biased kidney program. KidneyCellExplorer^19^ (https://cello.shinyapps.io/kidneycellexplorer/) shows growth hormone receptor is specifically expressed in male and female PT cells consistent with a growth hormone input. In addition, prolactin, the peptide hormone controlling postnatal functions such as milk production, is related to growth hormone and acts through its receptor (Prlr) to control Stat5a and Stat5b-directed transcription. *Prlr* shows one of the strongest biases in female enriched expression, consistent with prolactin signaling modulating female programs of kidney gene expression in association with reproduction. These observations argue for future studies focused on additional roles for growth hormone-Stat5a/Stat5b and prolactin-Stat5a/Stat5b regulation of female kidney programs.

Functionally, sex-biased genes are involved in multiple biological pathways, most notably peroxisomal lipid metabolism in the male and nuclear receptor pathways in the female. Proximal tubules utilize fatty acid as their major source of energy^86^, which is indispensable for their function in salt and water reabsorption. Peroxisomes oxidize long chain fatty acids while mitochondria break down short or medium sized fatty acids^87^. The bias for peroxisomal lipid metabolism possibly implies a higher energy demand in male proximal tubules than in female^88^, together with increased lipid deposition in the cortex^86^. In time of shortage in energy or oxygen (i.e., ischemia), it would be necessary to remodel renal expression profile towards a more energy-conserving state, which could explain the transient reversal of male phenotype during caloric restriction^63^ and short-term fasting^89^. Moreover, a byproduct of beta-oxidation is reactive oxygen species (ROS) produced by Acyl-CoA oxidases, which require antioxidant enzymes to neutralize. Excessive ROS due to high-energy state or reperfusion might contribute to chronic renal damage^90^. On the other hand, F-biased program highlight lipid clearance and anti-oxidation. F-biased genes *Abcc3* and *Gsta2/4/5* are all involved in the NRF2 pathway, whose activation acts against oxidative stress^91^ and inflammation^92^ to facilitate female resilience to kidney injury^93^.

AR-mediated gene expression is not only involved in the transport of organic anions (e.g., steroid conjugates and prostaglandins) and other solutes across the plasma membrane, but also directs cellular energetics and promotes lipid oxidation (see above) via Ppara and Nr1h4/FXR, both of which are nutrient-sensing TFs important for ciliogenesis in PTs^94^. This link between AR-dependent gene regulation and high-energy state to support renal function indicates how AR-mediated signaling could undermine renal function chronically, a potential cause for the M-biased susceptibility to kidney diseases. Interestingly, analysis of gene expression in published datasets of male mice undergoing 4-week caloric restriction^63^ highlighted a pronounced loss of AR-responsive gene expression and feminization of the male kidney. Caloric restriction has been reported to lower testosterone levels^95, 96^, which likely accounts for these observations and raises new questions about the complexity of actions on organ activity following an alteration of metabolism.

In the liver, hypothalamus-pituitary-directed pulsatile GH release dominate dimorphic gene expression in the liver^39, 41, 97, 98^ though our study shows 16% of M-biased genes in the liver were perturbed after adipocyte-specific removal of AR. The contrasting regulatory mechanisms of sexual dimorphism in the kidney and liver suggests that sexual differentiation of the two organs might have evolved separately and been selected for by different forces. Primarily functioning as a biochemical organ, the liver needs to respond to fluctuation in energy supply and to coordinate its enzymatic reactions to animal behaviors (e.g., food intake and physical activity)^99^. Direct link to the hypothalamus and pituitary can couple liver function to circadian/ultradian rhythms and fine-tune its action from hour to hour (a single GH pulse can alter Bcl6 expression^41^), as seen in the thyroid and adipose tissue^100^. In contrast, the functions of the kidney are fundamentally biophysical^101^ -- ultrafiltration and osmotic regulation^102^ -- where autoregulation is prevalent^103^. Androgens could reinforce the energetic profile of the kidney, but likely over a slightly longer time scale. In this regard, sexual differentiation of the kidney and liver plausibly allowed for adaptation to distinct environmental challenges during evolution.

Regarding renal sex differences in the human kidney, published analyses so far have not demonstrated extensive dimorphic expression, beyond the X and Y chromosomes, as reported here in the mouse PT^83, 85, 104, 105^. Limitations in variable sample quality and variation amongst individuals sampled is a confounding factor in the human studies. The full scope of human renal (and other organ) sex differences awaits further investigation. However, our re-analysis of human data revealed limited sex-dependent expression programs in the human kidney, with a modest conservation between human and mouse, lending support to a comparative approach^106, 107^ to study the molecular mechanism of sex differences in renal physiology and disease modeling.

## Acknowledgements

The authors thank members of the McMahon laboratory for helpful comments on experimental design and members of the Kim laboratory for useful discussion on single-cell analyses. Work in A.P.M.’s laboratory is supported by a grant from the National Institutes of Health (R01 DK126925). A.L.M. acknowledges support from the National Institutes of Health (R35GM143019) and the National Science Foundation (DMS2045327).

## Author Contributions

APM conceived the study. Funding support was generated by APM, JK and LP. Data were collected by JL, KK and JG and analyzed by LX, JL, SYH, MR, FG and IBH in consultation with APM, ALM, JK and LP. LX, JL, ALM and APM wrote the manuscript incorporating comments from all participants.

## Supplementary Figure Legends

**Figure S1. Transcript-level renal sex differences in adult mice.** Related to Fig. 1.

(A) Bar plot shows the fraction of sex-biased genes previously identified in the PT segments from our previous single-cell RNA-seq experiment^19^ (PTsex genes) that are covered in the core set of sex-biased genes identified by whole-kidney bulk RNA-seq experiment in the current study. Enrichment analysis was based on Chi-square test.

(B) Stacked bar plot shows the proportion of sex-biased transcripts identified in 8-week C57BL/6 mice that continue to show dimorphic expression in the aging kidney in the diversity outbred (DO) mice (the JAX data^45^) measured at 6, 12, and 18 months. The degree of overlap between 8-week C57BL/6 and JAX data is indicated by the progressive shading of the bars as age advances. The core set of sex-biased transcripts is defined as those that are shared by all four time points. Dashed boxes indicate the number and percentage of core sex-biased transcripts that are shared with PTsex genes.

(C) Read coverages of bulk RNA-seq at representative genomic regions demonstrate alternative splicing events.

(D) Read coverages of bulk RNA-seq at representative genomic regions demonstrate alternative isoform usage.

(E) Dot plot shows the enrichment of ToppFun pathways for both the core set of sex-biased genes and transcripts. Gene count of individual pathways is shown by the relative size of the dot, and the color of the dot indicates the statistical significance.

**Figure S2. Dynamical features of male and female renal transcriptomes during postnatal development.** Related to Fig. 2.

(A) Scaled average expression of core sex-biased genes over the five timepoints we measured, grouped by clusters shown in Fig. 2D. Together with the heatmap, we can see that neonatal genes are already expressed at high levels at early timepoints (0 and 2 weeks); pubertal genes exhibit increased expression over time (4 weeks onwards).

(B) Bar plot shows the number of core sex-biased genes in each cluster.

(C) Stacked bar plot indicates the fraction of neonatal and pubertal genes in the core set.

(D) Bar plot demonstrates the number of sex-biased transcripts identified at individual timepoints. Bars are colored by sex bias: M-biased genes are in green and F-biased genes in salmon.

(E) Heatmap shows the scaled average expression levels of the core sex-biased transcripts in male and female samples at individual timepoints. Genes were clustered via hierarchical clustering.

(F) Stacked bar plot indicates the fraction of neonatal and pubertal transcripts in the core set.

(G) Normalized read count for *Ar, Esr1* and *Hnf4a* over the five timepoints we measured. Lines indicate average expression levels. Boxes show the variation among samples. Data from male samples are in green and data from female samples are in salmon.

**Figure S3. AR-mediated signaling plays an important role in renal sex differences.** Related to Fig. 3.

(A) Bar plot shows the total number of differentially expressed genes identified in each treatment when compared to controls. Direction of perturbation in gene expression is indicated by the shade of the bar: up-regulated genes are in gray and down-regulated genes are in black.

(B) Scatter plots compare difference in the expression of core sex-biased genes in either of the two types of AR knockout experiments (nephron-specific removal [Six2-Ar-KO] and whole-body removal [Sox2-Ar-KO]) versus male-female differences at 8 weeks. Each dot represents a sex-biased gene, where sex bias is indicated by colors. Changes in gene expression (KO vs. WT or F vs. M) are summarized by log2 fold change (FC), which we used to evaluate the overall concordance of the two experiments by Spearman correlation.

(C) Three-way Venn diagrams for the M-biased genes that are common between treatments. Note that caloric restriction (CR) experiment^63^ on male mice perturbed almost half of AR-responsive M-biased genes in the kidney.

**Figure S4. Nephron-specific AR removal reverses dimorphic gene expression in males.** Related to Fig. 4.

(A) UMAP plot shows clustering outcome the RNA modality of the multimodal data (STAR Methods). Cluster nomenclature and color scheme are consistent with Fig. 4B.

(B) Dot plot of established marker expression for all cell populations.

(C) Bar plot shows the number of sex-biased genes identified in each PT segment.

(D) Bar plot shows the number of differentially expressed genes comparing M-KO to M-WT in each PT segment.

(E) Top bar plot shows the number of multimodal sex-biased genes and those identified from our previous single-cell RNA-seq experiment, with the number of overlapping sex-biased genes indicated on the right. Bottom bar plot shows the percentage of the overlapping sex-biased genes that were perturbed by nephron-specific AR removal. The number of genes that are perturbed in each treatment and the corresponding percentages are listed. Arrows indicate the direction of perturbation in gene expression as compared to controls.

(F) Volcano plots show multimodal sex-biased genes identified in PT-S1 and PT-S2 separately, where genes that are perturbed by AR removal in male are highlighted: comparing M-KO to M-WT, up-regulated genes are in red and down-regulated genes are in blue.

(G) Scatter plots contrast the effect of nephron-specific AR removal in male to the observed sex biases, among multimodal AR-responsive genes identified in PT-S1 and PT-S2, as shown in (E). Each dot represents a gene, where sex bias differs by color. Changes in gene expression (F-WT vs. M-WT or M-KO vs. M-WT) are summarized by log2 FC. Spearman’s rank correlation coefficient was calculated to assess the overall concordance of the two comparisons.

(H) Scatter plot compares the effect of nephron-specific AR removal in male to the observed sex biases, among multimodal AR-responsive genes identified in PT-S1, PT-S2, and PT-S3 combined. Each dot represents a gene, where sex bias differs by color. Changes in gene expression (F-WT vs. M-WT or M-KO vs. M-WT) are summarized by log2 FC. For genes that were perturbed in more than one segments, the maximum effect sizes are shown.

**Figure S5. Multimodal data reveals putative AR response elements near sex-biased genes.** Related to Fig. 5.

(A) Heatmap shows pairwise comparison of chromatin state between individual PT clusters. Scatter plot on the right exemplifies the correlation of normalized average peak counts for all PT marker peaks between PT-S3-mWT and PT-S3-f/mKO cluster.

(B) Bar plot shows number of significantly open regions for each condition of each pair-wise comparison (absolute Log2FC > 0.25, adjusted p-value < 0.05).

(C) Dot plot summarizes TF motif enrichment in the DARs within 100KB to TSS of the AR-responsive sex-biased genes in M-WT compared to F-WT PT segments.

(D) Volcano plot shows DARs within 100KB of sex-biased genes in PT-S1. 1,360 peaks were differentially open in male (left) and 1,273 peaks were differentially open in female (right). We colored F-biased peaks that are preferentially open in M-KO in red, and M-biased peaks that are preferentially closed in M-KO in blue, if any. Each dot represents a 500-bp region, where the nearest gene is annotated.

(E) Volcano plot shows DARs within 100KB of sex-biased genes in PT-S2. 6,444 peaks were differentially open in male (left) and 1,573 peaks were differentially open in female (right). We colored F-biased peaks that are preferentially open in M-KO in red, and M-biased peaks that are preferentially closed in M-KO in blue. Each dot represents a 500- bp region, where the nearest gene is annotated.

(F-K) Coverage plots of top sex-biased genes in M-WT, F-WT, and M-KO PT-S2/3: *Slco1a1, Cyp2j13, and Atp11a* (M-biased; F-H); *Hao2, Gsta4* (F-biased; I-K). Gene expression is also shown by violin plots adjacent to the tracks, with gene accessibility score *ψ* listed above each track. All peaks called in the region are shown in gray boxes, where DARs are highlighted in red. Peaks with potential AR binding site are indicated by red arrows.

**Figure S6. Fluorescent RNA in situ hybridization by RNAscope validates dimorphic gene expression in proximal tubule.** Related to Fig. 6.

RNAscope assay directly visualized the expression levels of sex-biased genes in M-WT, F-WT, and M-KO across the renal cortex. Scale bars = 100 μm.

(A) *Slco1a1* (red)*, Abcc3* (white)*, Cyp7b1* (green) co-stained with antibody against Aqp1 (blue).

(B) *Hao2* (red)*, Atp11a* (white)*, Cyp7b1* (green) co-stained with antibody against Aqp1 (blue).

(C) *Cyp2j13* (red)*, Cyp2e1* (Cyan)*, Cyp7b1* (green) co-stained with antibody against Aqp1 (blue).

(D) *Gsta4* (white)*, Cyp4a14* (red)*, Cyp7b1* (green) co-stained with antibody against Aqp1 (blue).

**Figure S7. Comparative study on kidney chromatin and hepatic gene expression.** Related to Fig. 7.

(A-B) Scatter plot compares changes in the expression of liver sex-biased genes that are in common between the in-house list and those identified from GSE112947 (A) and GSE174535 (B).

(C) Bar plot compares the number of differentially expressed genes identified in different treatment experiments.

(D) Scatter plot compares changes in the expression of liver sex-biased genes that are commonly perturbed by whole-body AR removal and castration in male mice.

## Supplementary Table Legends

**Table S1. List of sex-biased genes in the kidney identified from bulk RNA-seq data.**

**Table S2. List of differentially expressed genes in the kidney after various treatments.**

**Table S3. List of single-nuclear sex-biased genes in PT segments and the associated differentially accessible regions.**

**Table S4: Annotated DARs from pair-wise comparison among M-WT and M-KO vs F-WT PT segments within 100kb from TSS.**

**Table S5. List of sex-biased genes in the liver, and differentially expressed genes after systemic and hepatocyte-specific AR removal.**

**Table S6. List of sex-biased genes in human PT segments (re-analysis of GSE151302).**

## STAR Methods

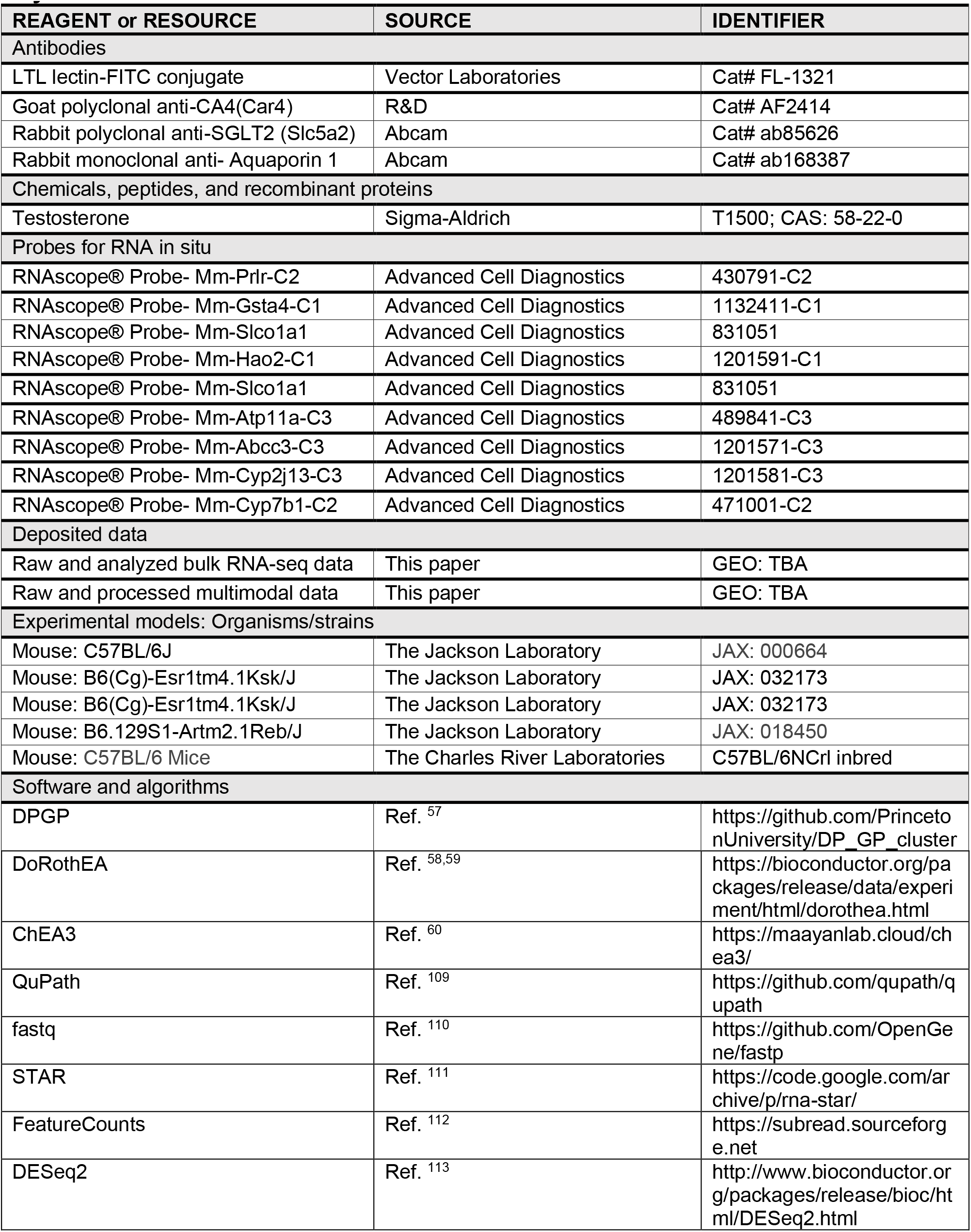
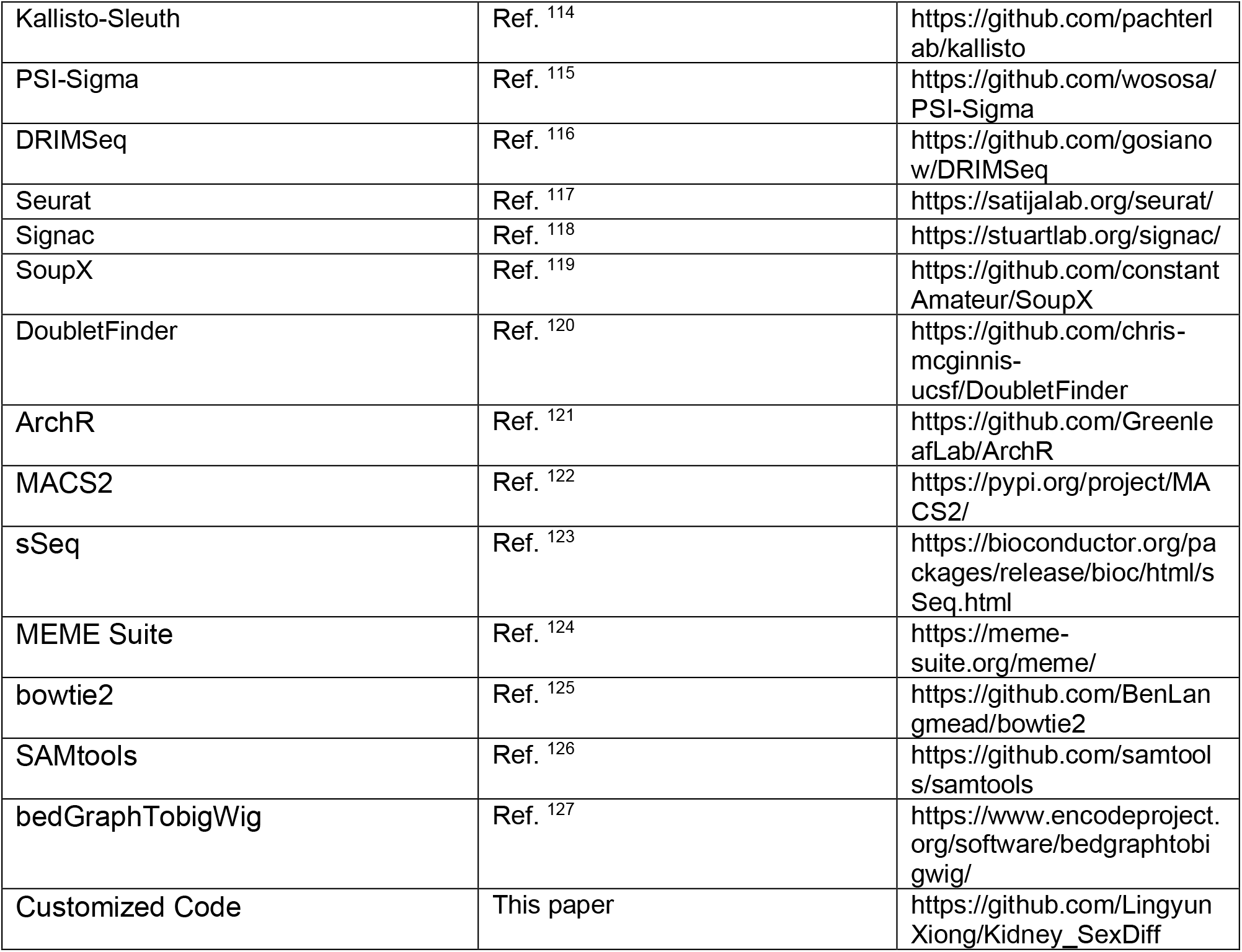
Key Resources Table.

### Lead Contact and Materials Availability

Further information and requests for resources and reagents should be directed to and will be fulfilled by the lead contact, Andrew P. McMahon. This study did not generate new unique reagents.

### Experimental Model and Subject Details

Institutional Animal Care and Use Committees (IACUC) at the University of Southern California reviewed and approved all animal work as performed in this study. All work adhered to institutional guidelines. Mice from the following strains were from the Jackson Laboratory: C57BL/6J (stock no. 000664), B6(Cg)-Esr1tm4.1Ksk/J (stock no. 032173), B6.129S1-Artm2.1Reb/J, (stock no. 018450), Six2TGC/+ mice were generated as described previously (Kobayashi A et a. 2008). Castrated males and ovariectomized females and control C57BL/6NCrl mice were from Charles River Laboratories.

### Method Details

#### RNA-seq

Whole kidney total RNA was extracted using Qiagen’s RNeasy Mini Kit and submitted to the Genome Access Technology Center at the McDonnell Genome Institute. Samples were prepared according to Clontech SMARTer library kit manufacturer’s protocol, indexed, pooled, and sequenced on Illumina HiSeq2500 platform for 50-bp single-end, and Illumina NovaSeq S4 for 150-bp pair-ended sequencing.

#### Single-nucleus multimodal experiment

C57-B6J adult mice (age: 9-12 weeks old; sex: males weighing 21.5-23.2 grams and females weighing 18.4-19.4 grams) were euthanized with CO2 chamber and perfused with ice cold HPBS (Hyclone). The kidney capsules were removed, and kidneys cut into 6 smaller pieces and flash frozen in liquid nitrogen for further processing. On the day of, nuclei were isolated as previously described^128^. Briefly, flash frozen kidney pieces were thawed on ice and minced into small pieces (<1mm) with a sterile razor blade and then dounced in Nuclei EZ Lysis Buffer (Sigma) with Protease Inhibitor. The tissue was dounced 15X loose, filtered with a 200uM filter, and then 5X tight, incubated for 5 mins on ice, filtered through 40uM filter and spun down at 500G x 5min at 4°C in a swinging bucket centrifuge. Supernatant was removed and nuclei pellet resuspended in Nuclei EZ lysis buffer, incubated an additional 5 mins on ice and spin down again. The final pellet was resuspended in Diluted Nuclei Buffer provided in the 10X Chromium Kit and passed through a pre-wetted 5uM filter. All buffers had 1U/ul Protector Rnase inhibitor and 1mM DTT added to preserve RNA integrity. Nuclei were then counted on a Countess III machine and the targeted number (∼9,000 nuclei/sample) was loaded into a GEM J Chip as per manufacturer’s specifications. Multiomic (10x – PN:10002805) reagents and index plates were used to generate the snATACSeq and snRNASeq libraries. Libraries were processed using 10X Genomics Manual CG000338 (7 preAmp, 8 ATAC library construction, 8 cDNA, 16 GTEX Sample Index – PCR cycles) Libraries were checked by BioAnalyzer before sending to Novogene for NovaSeq6000 S4 PE150 sequencing using Illumina platform.

#### Fluorescence RNA in situ hybridization

RNA In situ hybridizations were performed following RNAscope Multiplex Fluorescent Reagent Kit v2 user manual (Advanced Cell Diagnostics) as previously described^19^. We used antibodies for immunofluorescent co-staining on the same frozen sections with RNA probes: LTL lectin-FITC conjugate (#FL-1321; Vector Laboratories); Aqp1 (# ab168387, rabbit; Abcam), SGLT2 (Slc5a2) (# ab85626, rabbit; Abcam), Car4 (#AF2414, goat; R&D). Number of subcellular dots from RNAscope experiments were quantified through QuPath^109^. Briefly, cells were detected by nucleus staining, then only the spots of target gene with positive PT-segment marker expression were counted.

### Quantification and Statistical Analysis

#### Bulk RNA-seq Data Analysis

Bulk RNA-seq data generated in this study and public data (kidney: GSE121330; liver: GSE112947 and GSE174535) were analyzed using a custom workflow, which is available on GitHub and briefly described in the following. First, raw sequence reads were pre-processed using *fastp*^110^ (version 0.23.2), which was used to trim low quality (quality score <=20) and to filter short reads (<=20bp). Sequence reads passing quality control were aligned to mouse genome build mm39 (GRCm39) using *STAR*^111^ (version 2.7.0) and those that mapped to annotated genomic regions (GENCODE release M28) were counted using *FeatureCounts*^112^ (version 2.0.3). Differential expression analysis was performed using *DESeq2*^113^. Genes with low count per million values (CPM<1) were excluded, and differential expressed genes were identified based on thresholds of adjusted *P-value* (<0.05) and absolute log2 fold change (>0.5; i.e., greater than 1.41- fold). Overall sample variation was evaluated by principal component analysis implemented in *DESeq2*, and batch effect (if present) was accounted for by specifying batch information as a covariate in the regression model. Average TMM-normalized gene expression^129^ was used as input for heatmap visualization, and scaled expression levels across developmental timepoints or treatment conditions were shown.

#### Isoform analysis

Isoform expression of known Ensembl transcripts were estimated with Kallisto^114^. For all the analyses, only transcripts with (a) adjusted *P*-values < 0.05, (b) absolute log2 fold change > 0.5, and (c) TPM > 1 were kept. Alternative splicing analysis was performed using PSI-Sigma^115^ with default settings. Differential transcript usage was identified using DRIMSeq^116^. A3SS: alternative 3’ splice site; A5SS: alternative 5’ splice site; IR: intron retention; MES: multi-exon skipping; SES: single exon skipping; TSS: transcription start site.

#### Functional Inference

Pathway enrichment analysis of sex-biased genes was performed using *ToppCluster* web browser^49^ with default settings. We used *GOATOOLS*^130^ for gene ontology analysis, where only terms of biological processes were considered, and multiple testing correction was performed with the Benjamini-Hochberg method.

#### Computational prediction of upstream regulators

Temporal co-regulation of sex-biased genes was studied using *DPGP*^57^, where difference in average TMM-normalized gene expression between male and female samples over 5 timepoints were clustered based on shared dynamical features. Clusters with the most genes were prioritized for visualization. Scaled expression levels were shown. The *DoRothEA*^58, 59^ and *ChEA3* web browser^60^ were used to infer upstream regulators of sex-biased programs in the kidney. The input for *DoRothEA* was TMM-normalized expression of sex-biased genes or known proximal tubule markers; only high-confidence regulons (class A and B) were used to compute normalized enrichment score (NES) of curated TFs. Given the M-or F-biased genes, *ChEA3* ranked TFs by weighing and integrating extensive ChIP-seq and co-expression evidence for putative TF-target relationship. Mean rank was used in this study. The top 15 TFs were selected to view co-expression networks, whose expression levels in proximal tubule were checked against previous scRNA-seq data^19^ (https://cello.shinyapps.io/kidneycellexplorer/). Putative target genes of top-ranked TFs predicted by *ChEA3* were examined for relative impact and overlaps.

#### Single-nuclear multimodal data processing

We used *Seurat*^117^ (version 4.1.0) and *Signac*^118^ (version 1.5.0) in R (version 4.0) for primary multimodal (RNA and ATAC) data processing, following the guidelines provided by the software developers. Briefly, we loaded both modalities for each sample and merged counts into a single data object for general data quality evaluation. We assessed ambient RNA contamination in each sample using *SoupX*^119^, to find that the global contamination was 2-4%. Doublets were detected using *DoubletFinder*^120^, and were filtered together with low-quality cells by the following cut-offs: RNA feature (250-7,000), percentage of mitochondrial RNA (<35%), total ATAC count (1,000-100,000), nucleosome signal (<2) and transcription start site (TSS) enrichment score (>1).

RNA-data were log-normalized and scaled based on the top 2,000 variable features and the data were projected to lower dimension using principal component analysis (PCA). After evaluation of elbow and jackstraw plots, the top 30 principal components were used for k-nearest neighbor (kNN)-based clustering, with a resolution of 0.5. Considering known biological differences between male and female kidneys, we pooled samples using reciprocal PCA-based integration method implemented in *Seurat*. ATAC-data were processed using performing widely implemented latent semantic indexing (LSI) method. We performed term frequency-inverse document frequency (TF-IDF) normalization, followed by top feature identification and singular value decomposition. Leveraging information from both data modalities, the joint neighbor graph was constructed for final clustering using the weighted nearest neighbor methods implemented in *Seurat*. Clustering outcomes were visualized in UMAP plots, where depth imbalance was noted between male and female samples. Features enriched for individual clusters were identified by Wilcoxon rank-sum test with a cutoff for minimum log2 fold change (>0.25) and minimum percentage of cells with expression (>0.25). For cell type annotation, top features of each cluster were compared against established markers for broad cell types known to be present in the kidney^19, 131^. Normalized gene expression data were visualized in feature, dot, and violin plots; normalized peak counts were visualized in coverage plots with peaks highlighted.

We used *ArchR*^121^ (version 1.0.1) for additional multimodal data processing. Besides standard filtering criteria as above and iterative LSI dimensionality reduction with default settings, we created pseudo-bulk replicates for each cluster and performed customized peak calling using *MACS2*^122^ with an FDR cutoff of 0.05. Besides standard genomic features, the peak matrix was also annotated with canonical TF motif using the motif set curated by the Jasper 2022 CORE database^108^ (Mus musculus) for AR (MA0007.3) and Hnf4a (MA0114.3), as well as publicly available ChIP-seq data for AR and Hnf4a (see below). The peak matrix was then categorized by the distance to the nearest TSS: peaks within 1 kb of TSS were categorized as proximal peaks; peaks within 100kb but not within 1 kb of TSS were categorized as distal peaks.

#### Single-nucleus multimodal differential analysis

To identify differentially expressed genes between two annotated clusters of interest, we performed proportional fitting-based depth normalization on raw read counts to mitigate depth imbalance, before applying *sSeq*^123^ as described previously^19^, with a cutoff of adjusted *P-*value < 0.05 and absolute log2 fold change > 0.5 (i.e., greater than 1.44-fold). *sSeq* is a shrinkage-based method for estimating dispersion in negative binomial models for RNA-seq data, well suited for small sample sizes. Briefly, we treated annotated clusters as meta-cells, and computed average normalized gene expression and proportion of non-zero expression cells for each gene across meta-cells. We identified differentially expressed genes for each PT segment separately. The number of differentially expressed genes recovered at meta-cells was significantly higher than those detected at single-cell level, and the signals were shown to be more robust and comprehensive^19^.

We identified differentially accessible regions (DARs) between two clusters of interest using Wilcoxon test, adjusted for TSS enrichment score and number of unique fragments per cell, with a cutoff of FDR < 0.05 and absolute log2 fold change > 0.25. DAR identification was performed for each segment of the PT cell types. Intersections of the pairwise DARs were categorized and visualized with upset function in *UpSetR* package. To identify DARs that are differentially open in the male WT compared to female WT and male KO, we found the intersection of male WT vs female WT DARs and male WT vs male KO DARs, each with positive log2 fold-change value for male WT. To identify DARS that are differentially closed in the male WT compared to female WT and male KO, we found the intersection of male WT vs female WT DARs and male WT vs male KO DARs, this time with negative log2 fold-change value for the male WT. DARs were examined for nearest genomic features and TF motif/binding enrichment, based on original peak annotations specified above.

#### Gene accessibility score

The gene accessibility score *ψ* is a metric that quantifies how open a genomic region is by summing peak access within a gene body and some distance upstream of its transcription start site (TSS), weighted by the distance of the peak to the TSS and the variability in peak accessibility across cell types, as defined by Janssens et al.^132^. In this study, we used normalized accessibility of each peak per cell as the input, including all peaks inside the gene body and up to 5kb upstream of TSS, but excluding peaks residing within the body of nearby genes. The gene accessibility score was computed as the weighted sum of individual peak accessibility, where the total weight for each peak is the product of the distance and the variation weight. The distance weight is assigned to each peak using an exponentially decaying function so that peaks further away from the TSS are given lower priority, as implemented by the function of calculating gene score in *ArchR*. To prioritize peaks with variable or differential accessibility across cell types, we calculate the Gini coefficient of each peak among all clusters and use the z-normalized Gini coefficient as the exponent for the variation weight. For visualizing accessibility of selected genes in the heatmap (**Fig. 6D**), the gene accessibility score calculated for each gene was scaled by the maximal value across clusters.

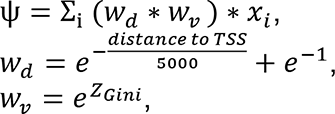

where *x_i_* was the normalized peak count.

#### Motif Analysis

Matching of canonical TF motifs was performed using the Find Individual Motif Occurrences (FIMO) function in MEME Suite^124^ (version 5.5.0), where we supplied PWM information for AR (MA0007.3) and Hnf4a (MA0114.3) from the Jasper 2022 CORE database^108^ (Mus musculus). We identified annotated TF motifs that are enriched among peak sets of interest using the Simple Enrichment Analysis (SEA) function in MEME Suite^124^ (version 5.5.0), by specifying the motif database to be HOCOMOCO mouse (v11 full)^133^. We used a *p*-value cut-off of 1E-5 for motif matching and enrichment analysis. Either random genomic regions with matching GC-content or shuffled input sequences were used as the background for comparison. Multiple-hypothesis testing correction was performed using the Benjamini-Hochberg method.

#### ChIP-seq data processing

Public ChIP-seq data for AR and Hnf4a binding sites in adult kidney tissues (GSE47194^46^) were processed through a custom pipeline developed in the lab. First, raw sequence reads were pre-processed using *fastp* (version 0.23.2; Chen et al., 2018), during which reads were trimmed and reads of low quality (quality score <= 20) and short length (<= 20bp) were filtered. Sequence reads passing quality control were aligned to mouse genome build mm10 (GRCm38) using *bowtie2*^125^ (version 2.3.5) and *SAMtools*^126^ (version 1.10). Peaks mapped to annotated genomic regions (GENCODE release M22) were called for TF-treated bam files against controls using *MACS2*^122^ with an FDR cutoff of 0.05, specifying no lambda or model, and allowing for a shift size of 75 base pairs and extension size of 150 base pairs. If ChIP-seq experiment was repeated (such as AR-ChIP-seq in GSE47194), replicated peaks were defined as TF-binding sites. In the multimodal data, all accessible peaks overlapping with TF-binding sites with a maximum gap of 250 base pairs were annotated as TF-bound. We used *bedGraphTobigWig*^127^ software (version 2.8) to convert the peak files to bigwig files for genome browser visualization.

### Data and Code Availability

- RNA-seq data have been deposited at Gene Expression Omnibus and are publicly available as of the date of publication.
- All original code has been deposited on GitHub (https://github.com/LingyunXiong/Kidney_SexDiff).
- Any additional information required to reanalyze the data reported in this paper is available from the lead contact upon request.

