## Supplementary figures and images for "Direct androgen receptor regulation of sexually dimorphic gene expression in the mammalian kidney"

### SI Figures

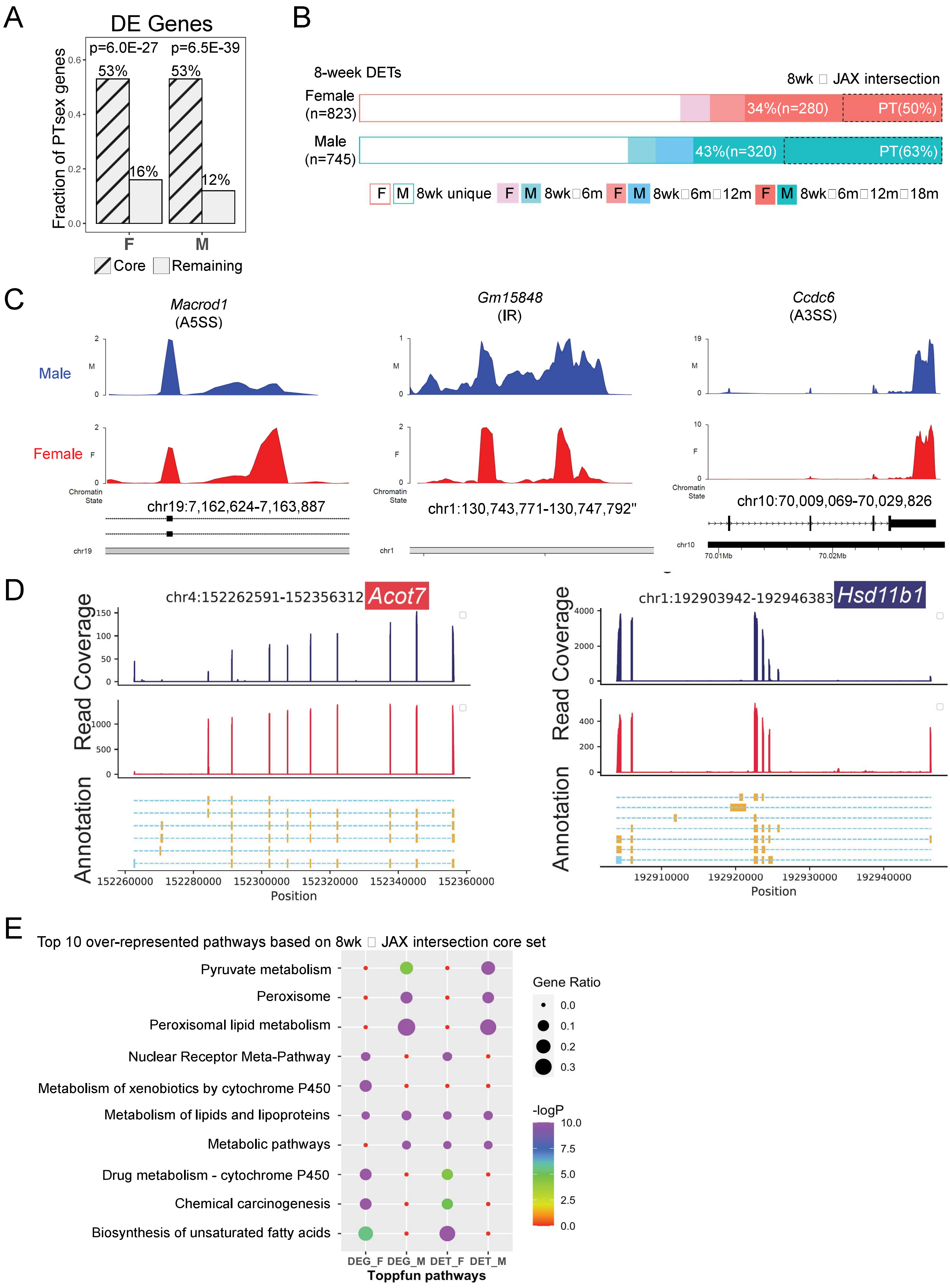

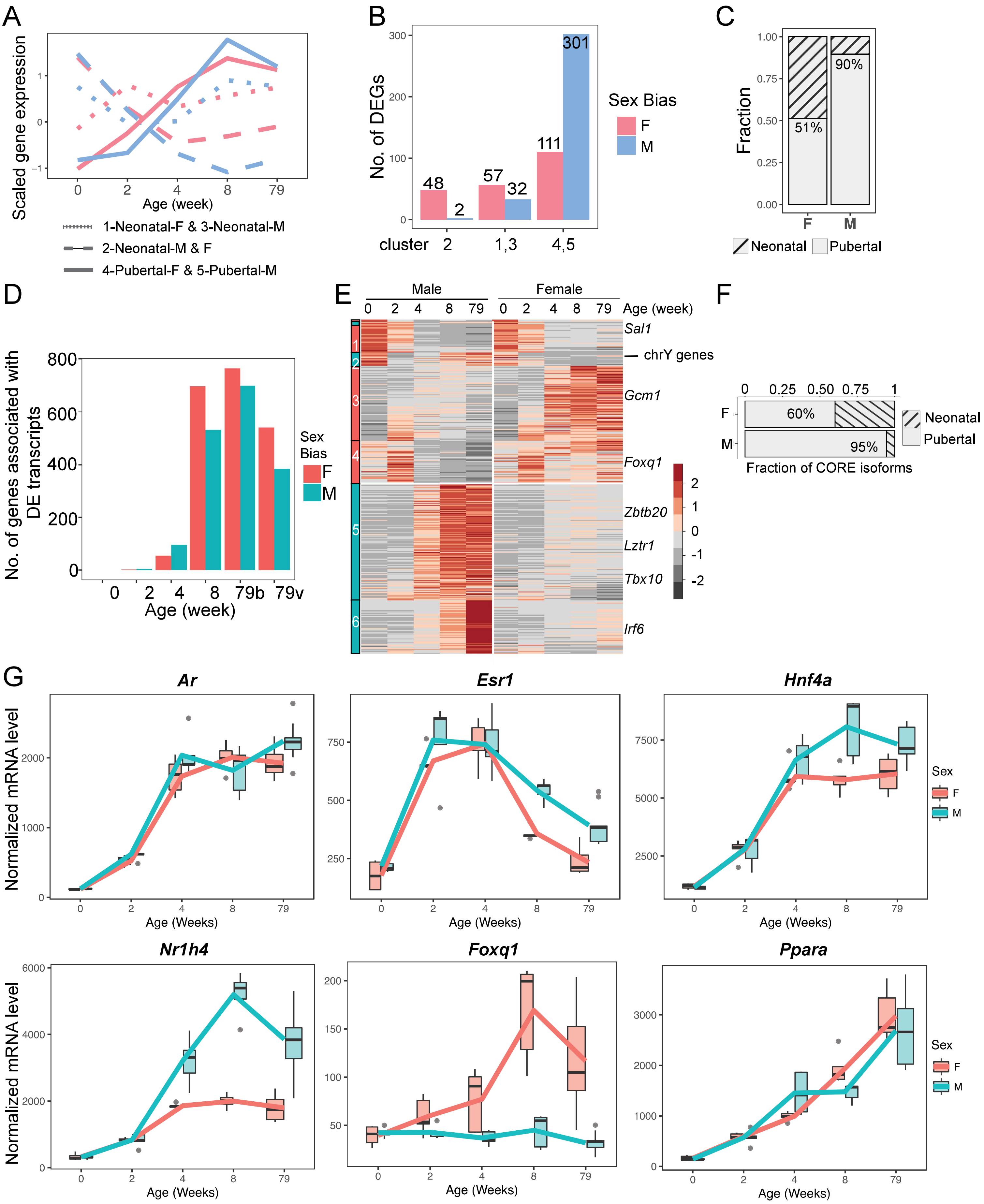

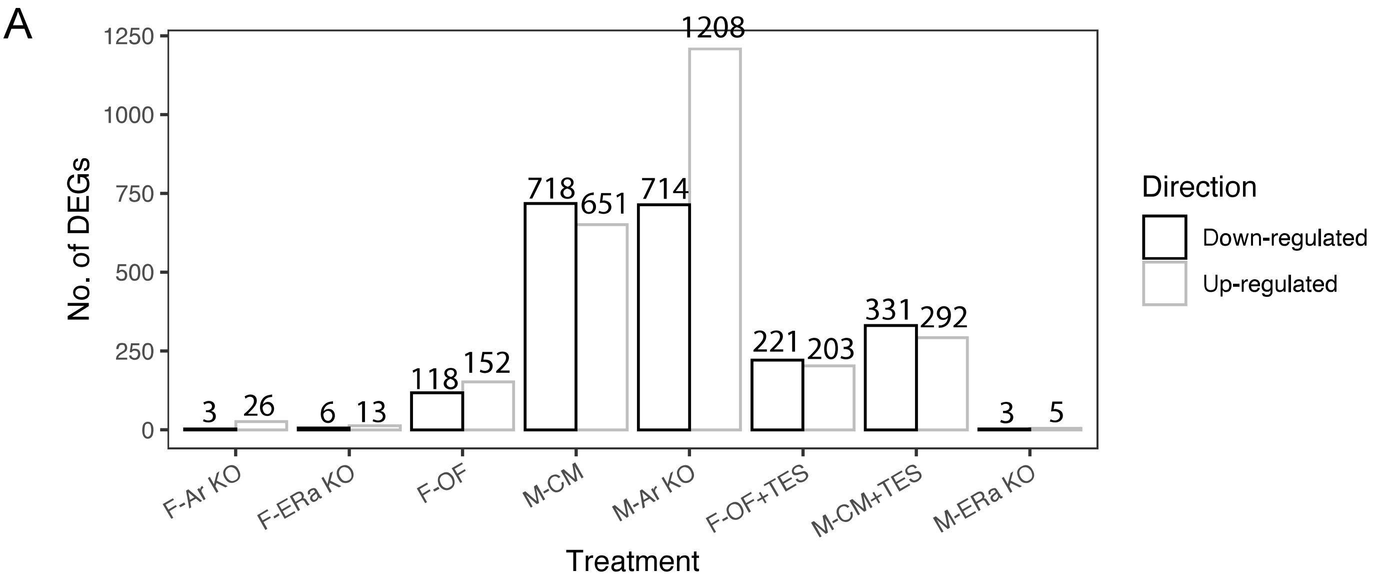

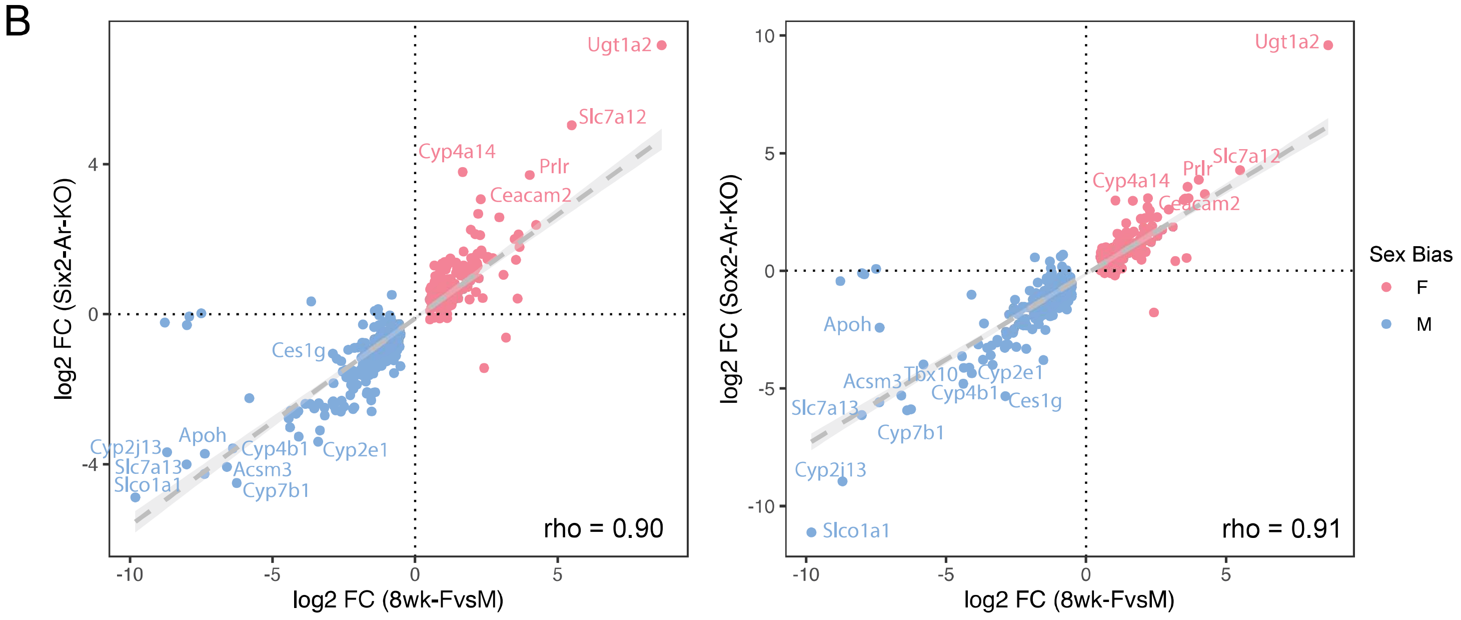

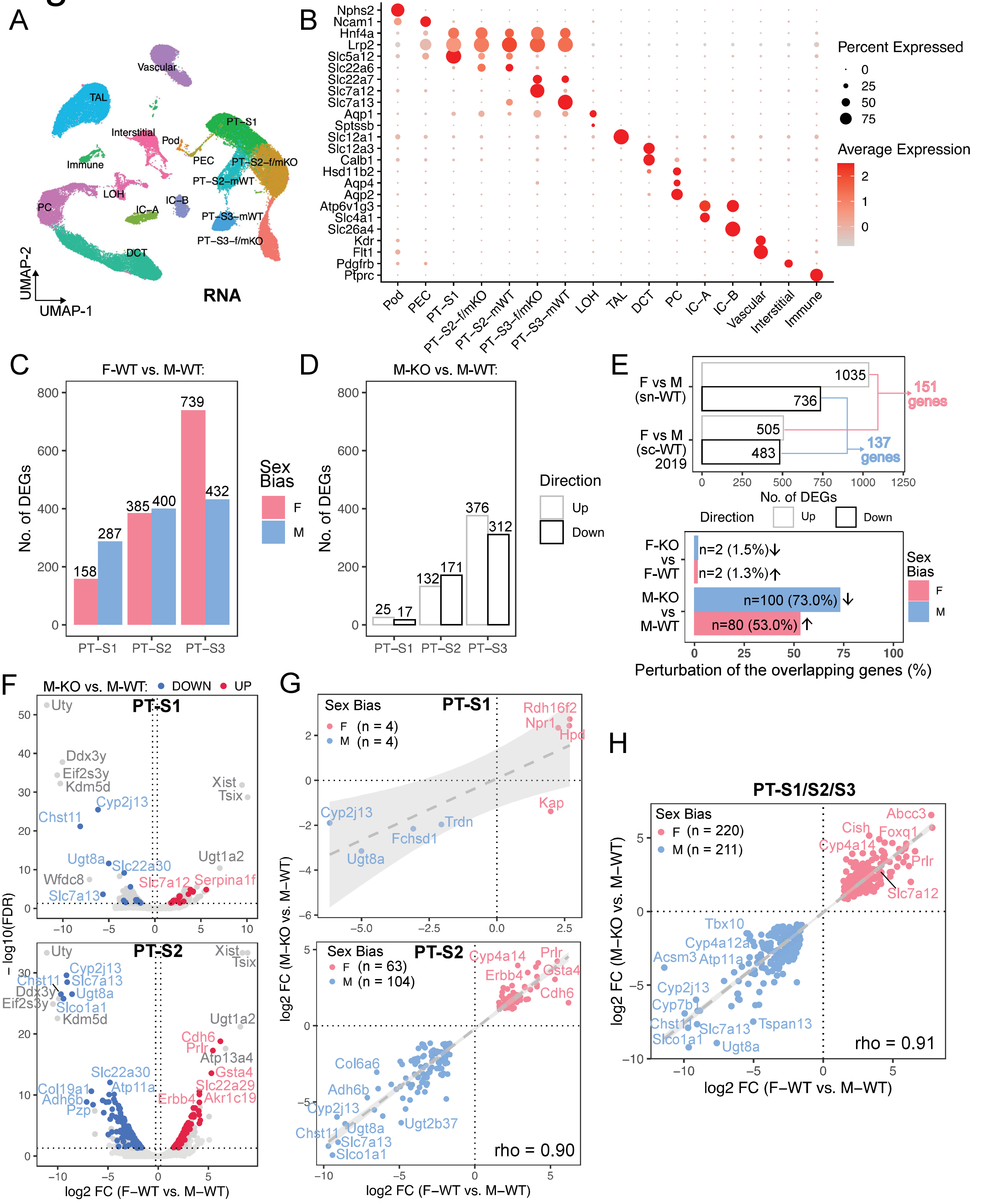

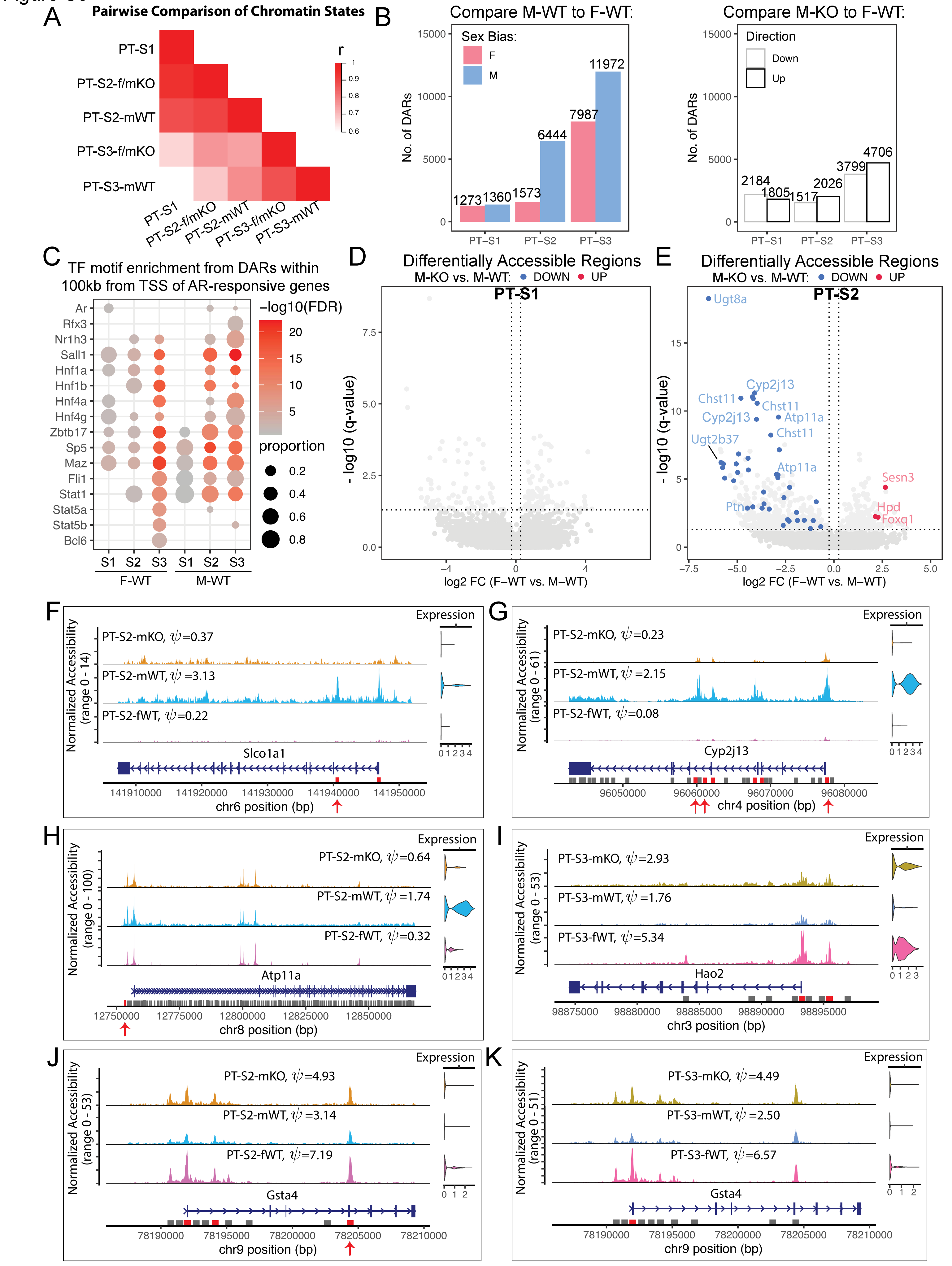

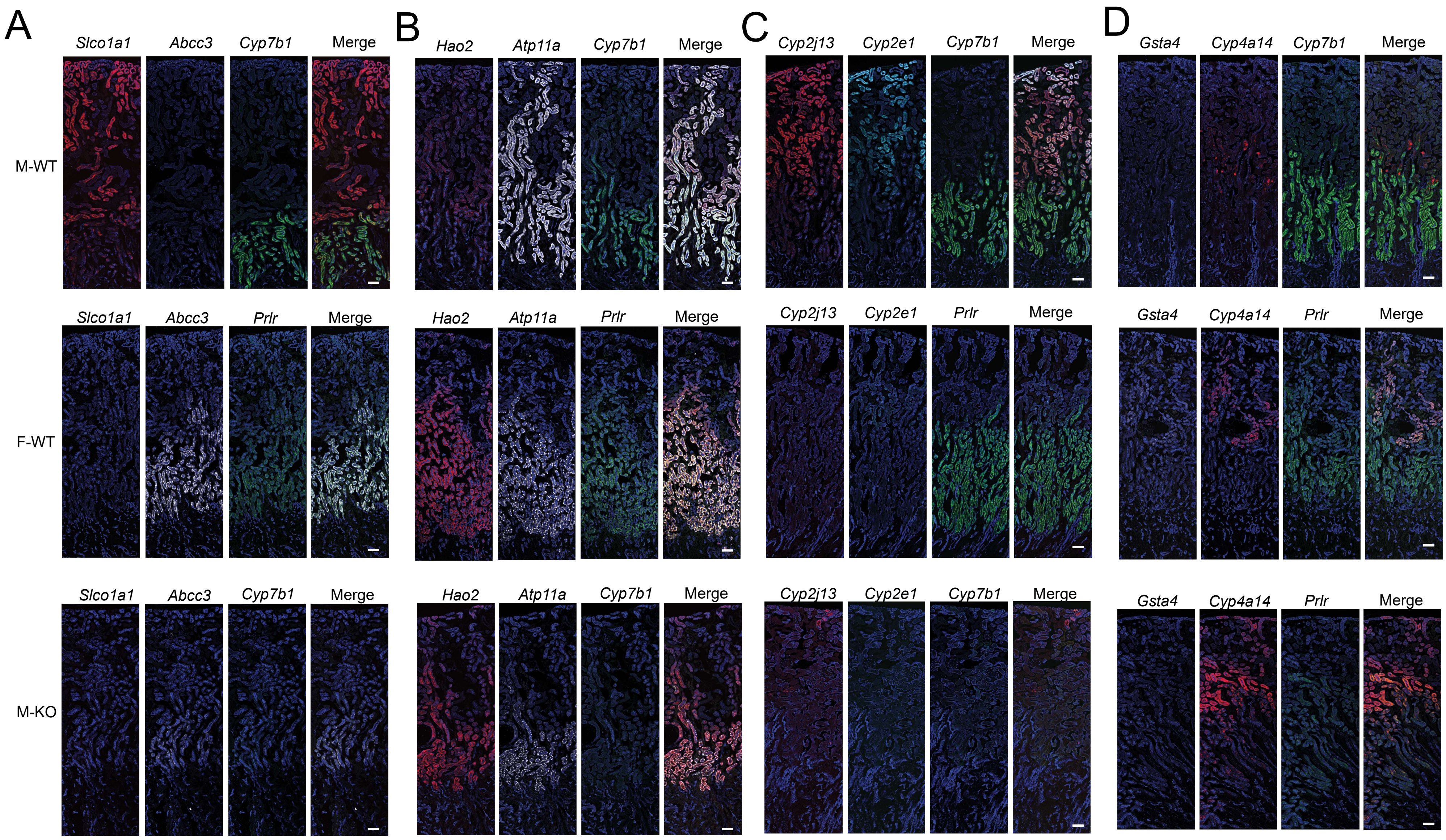

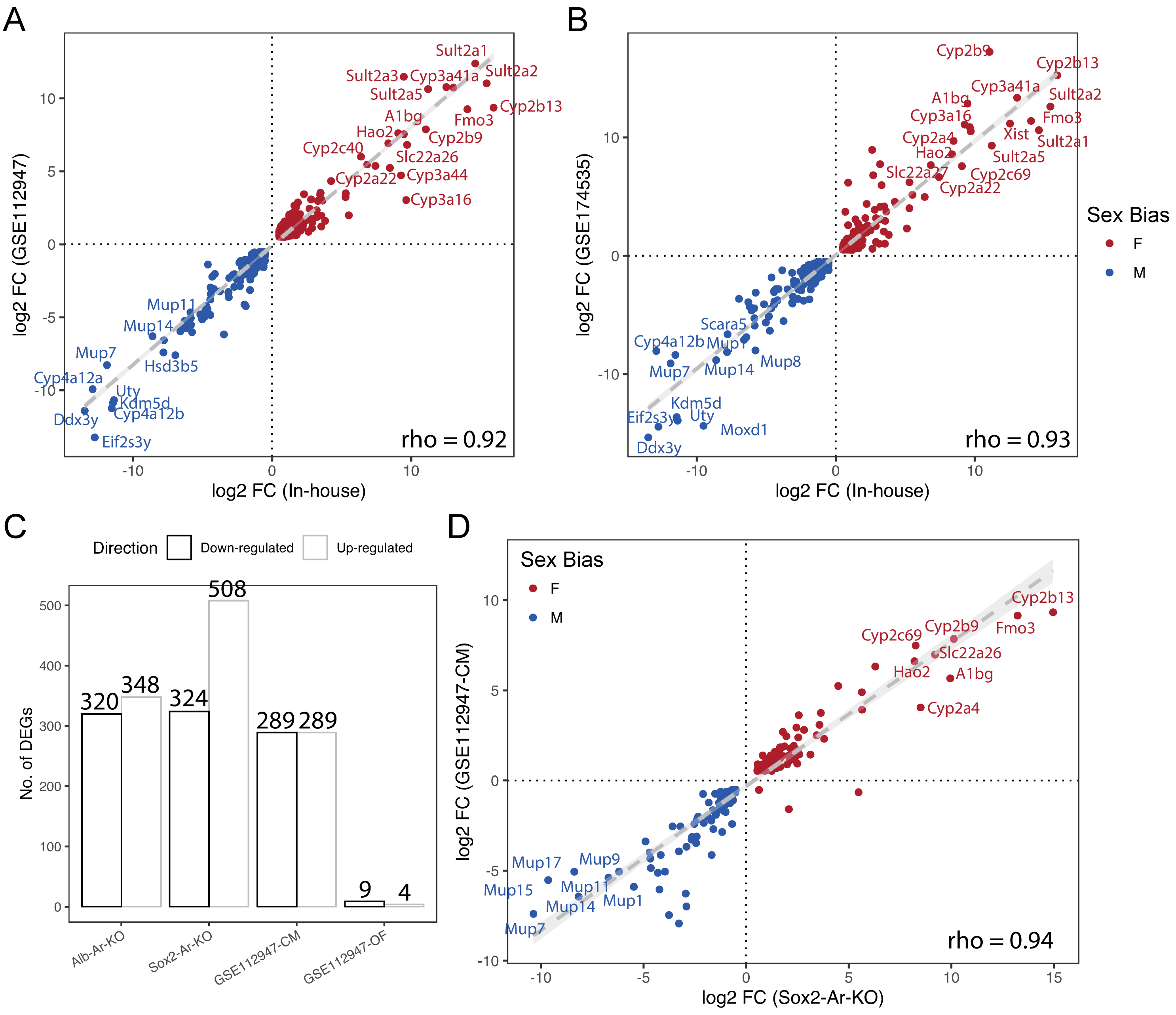
